# Tumor microenvironment distinctions between esophageal cancer subtypes explain varied immunotherapy responses

**DOI:** 10.1101/2024.09.24.614705

**Authors:** Seungbyn Baek, Junha Cha, Min-Hee Hong, Gamin Kim, Yoon Woo Koh, Dahee Kim, Martin Hemberg, Seong Yong Park, Hye Ryun Kim, Insuk Lee

**Author notes:** Corresponding authors: Seong Yong Park, Hye Ryun Kim, Insuk Lee.

## Abstract

Esophageal cancer comprises two main subtypes: esophageal squamous cell carcinoma (ESCC) and esophageal adenocarcinoma (EAC). Previous studies have revealed distinct genomic characteristics between these subtypes, with ESCC sharing similarities with head and neck squamous cell carcinoma (HNSCC), and EAC aligning with gastric adenocarcinoma (GAC). Additionally, recent immunotherapy clinical trials have shown a higher response rate in ESCC and HNSCC compared to EAC and GAC. However, how the tumor microenvironment contributes to these varied immunotherapy responses remains unclear. In this study, through a comparative analysis of single-cell tumor transcriptomes from 35 patients with ESCC, EAC, HNSCC, or GAC, we identified two groups with tumor microenvironment distinctions: ESCC and HNSCC versus EAC and GAC, consistent with their genomic classifications. Malignant epithelial cells displayed distinct separations based on histological origin. In the tumor immune microenvironment, we observed an enrichment of CXCL13^+^CD8^+^ T cells and CXCL9^+^CXCL10^+^ tumor-associated macrophages (TAMs) in ESCC and HNSCC, which activate cellular immunity through interferon-γ. In contrast, EAC and GAC exhibited a high presence of heat-shock protein-expressing CD8^+^ T cells and MARCO^+^ TAMs. These immune signatures help explain the varied immunotherapy responses among these cancer subtypes and successfully predict immunotherapy outcomes across diverse cancer types, underscoring their clinical significance.

## Introduction

Esophageal cancer is one of the most lethal cancers, ranking as the 7th most common cause of cancer death worldwide, with a 5-year relative survival rate of 21% in the United States^1^. Esophageal cancer consists of two anatomically shared but histologically different types: esophageal squamous cell carcinoma (ESCC) and esophageal adenocarcinoma (EAC). ESCC is more prevalent in East Asia, the Middle East, and East and South Africa, with main risk factors including smoking, alcohol consumption, and regional micronutrient deficiencies^2^. Conversely, EAC is more prevalent in North America, Europe, and Australia, with obesity and acid or bile reflux being major risk factors. Low-grade squamous hyperplasia and high-grade dysplasia develop into carcinoma in ESCC, whereas the pre-malignant metaplastic epithelium known as Barrett’s esophagus eventually transforms into EAC through dysplasia^2,3^.

Several studies have compared the various omics profiles of ESCC and EAC. Genomic profiling analyses have revealed that while both ESCC and EAC show a high frequency of genomic alterations, they differ in their profiles^4^. Molecular and genomic profiling conducted by The Cancer Genome Atlas (TCGA) program revealed distinct genomic characteristics for ESCC and EAC, while also highlighting similarities with nearby cancer types^5^. ESCC is more related to the ‘classical’ subtype of head-and-neck squamous cell carcinoma (HNSCC) without HPV infection. In contrast, EAC often shares anatomical regions with gastric adenocarcinoma (GAC) at the gastro-esophageal junction (GEJ) and exhibits profiles similar to GAC characterized by chromatin instability. Therefore, an in-depth investigation of the cellular and molecular distinctions between these groups of nearby cancer types would facilitate our understanding of the varying clinical outcomes between ESCC and EAC.

Recently, the application of immune checkpoint blockade (ICB) for esophageal cancer has been widely studied. Due to their locational similarities, various clinical trials have been conducted for patients with advanced/metastatic ESCC or EAC^6–8^. By directly comparing responses to anti-PD-1 treatment, two out of three clinical trials showed better response rates for ESCC patients, while the other trial reported similar responses for both ESCC and EAC patients. Furthermore, recent clinical trials with anti-PD-1 and anti-CTLA-4 combination therapy showed improved efficacy for ESCC^9^, partial and limited responses for HNSCC^10^, and almost no improvement compared to chemotherapy for GAC and EAC at the GEJ^11^.

Since the tumor microenvironment (TME) influences ICB responses, we hypothesized that distinct intra-tumor immune cell populations might contribute to the different response rates for ICB treatment. In the present study, to make cellular differences between ESCC and EAC more pronounced and identifiable, we conducted single-cell gene expression analysis on tumors from 35 patients with four cancer types: ESCC, EAC, HNSCC, and GAC, all located near the esophagus. Through the comparison of these four cancer types with resolved cellular heterogeneity, we identified distinct cellular components between the two subtypes of esophageal cancer and their shared similarities with nearby cancer types in the TME, which can explain their different responses and predict ICB responses across diverse cancer types (**Figure 1a**).

**Fig. 1.**
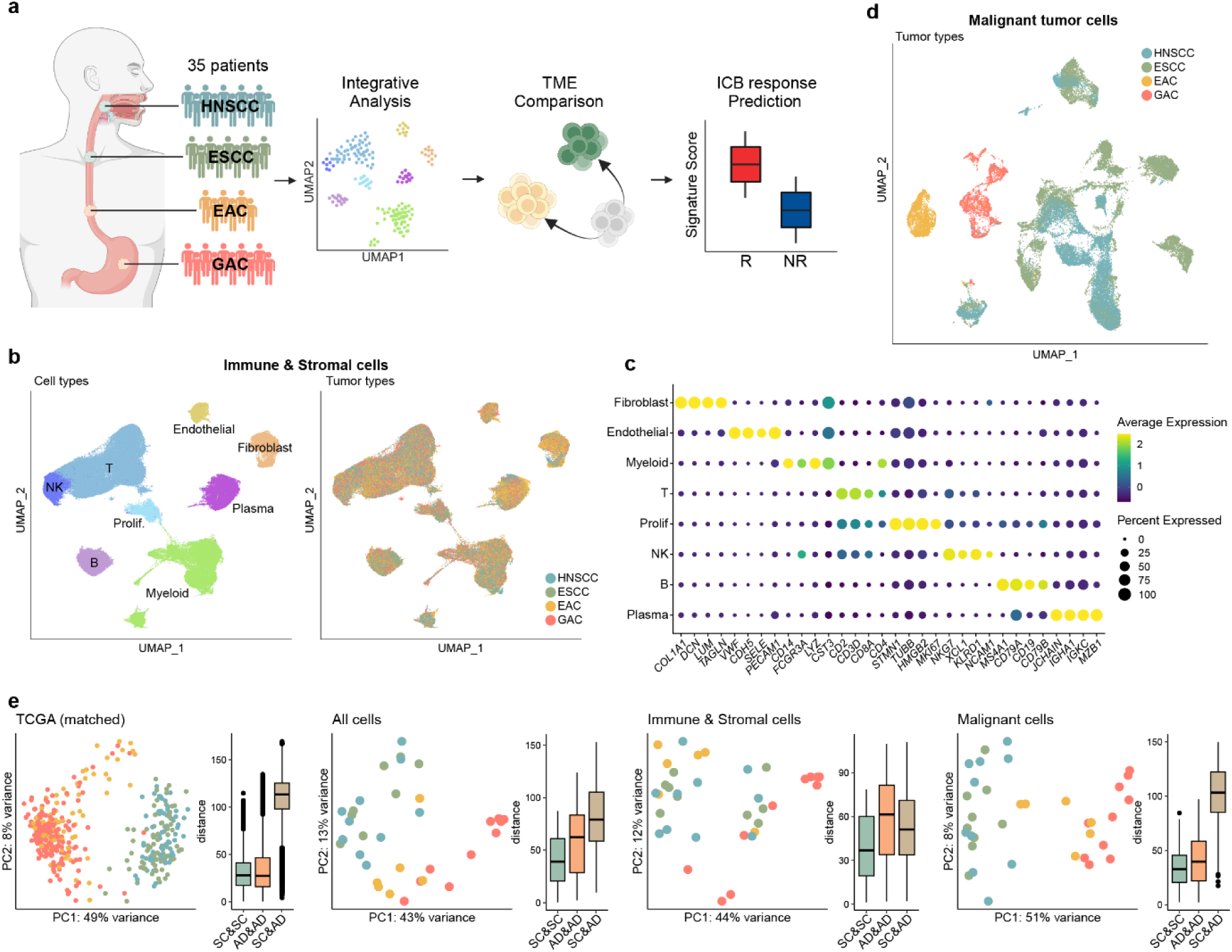
Study overview. **a**, Study scheme. **b**, UMAP plot of integrated immune and stromal cells from all cohorts combined colored for (Left) cell types and (Right) tumor types. **c**, Dotplot of markers genes for each major cell type. The colors indicate scaled average expression for each subpopulation and the sizes indicate percentage of cells that are expressing each gene. **d**, UMAP plot of integrated malignant cells from all cohorts colored for tumor types. **e**, PCA plots for samples and boxplots for pair-wise distances among the samples grouped by combinations of histological origins of tumor cells.

## Results

### Integration of single-cell transcriptome for four cancer types near the esophagus

For the analysis of four cancer types near the esophagus at single-cell resolution, we generated single-cell transcriptome datasets from patients with HNSCC, ESCC, or EAC without previous history of treatment. Furthermore, we collected additional single-cell EAC^12^ and GAC^13^ datasets that were generated using 10X Genomics platform and had comparable data quality and cell counts to the in-house datasets. To ensure maximal hypothetical similarity to patient conditions as characterized in the TCGA study on esophageal cancer^5^, we filtered the HNSCC and GAC datasets accordingly. For the HNSCC dataset, we selected samples from patients without HPV infection and excluded samples collected from the oral cavity or tonsillar regions. For the GAC dataset, we chose samples from patients annotated as having chromatin instability (CIN) subtype from the original article (**Supplementary Table 1-2**). We carefully tested and selected combinations of preprocessing steps for accurate integration of these datasets (**Supplementary Figure 1a-e**). Through thorough integration, we compiled a total of 150,405 immune and stromal cells (**Figure 1b**) and annotated the major cell types with their markers (**Figure 1c, Supplementary Table 3**). Additionally, we integrated 53,353 epithelial cells, of which 40,959 were malignant tumor cells (**Figure 1d**).

Because bulk-level transcriptome profiles for the four cancers showed clear distinction based on their histological origins of epithelial cells (i.e. squamous epithelial cells for squamous cell carcinomas and glandular epithelial cells for adenocarcinomas) from TCGA analysis^5^, we wanted to determine if these distinct patterns could be observed in transcriptomes from specific cell types. We compared distinctions among the four cancers using bulk transcriptome datasets from TCGA and pseudo-bulk samples generated with the single-cell datasets, based on different cell type criteria. We visualized each sample on PCA coordinates to analyze their distributions based on similarity and calculated pair-wise distances between samples (**Figure 1e**). Pseudo-bulk samples with only malignant tumor cells showed clear distinction and close distances among samples with the same histological origins, as observed in TCGA bulk analysis. This is because malignant cells have different histological origins, and bulk transcriptome datasets tend to consist of 60∼90% of epithelial cells due to tumor purity thresholds^14^. Pseudo-bulk samples generated with all cell types still showed distinctions by histological origins, but to a less degree compared to those from malignant cancer cells. However, we observed less separations for pseudo-bulk samples composed of immune and stromal cells. We hypothesized that while non-malignant cells in the TME have differences in underlying immune mechanisms for each cancer, they tend to have more shared apparent features and similarities compared to the more distinct epithelial cell types. With this preliminary analysis, we aimed to further explore distinctions in malignant cells regarding cell states and functions and to dissect immune mechanisms within immune and stromal compartments that could differentiate each cancer type.

### Distinction of malignant cells between ESCC and EAC by histological origins

Since we identified the most coherent distinctions among malignant tumor cells, we analyzed similarities among tumor types and patients using a single-cell resolution approach. We applied differential expression gene (DEG) analysis comparing malignant cells from ESCC and EAC patients, which revealed clear separation between these DEGs based on their epithelial cell origin (**Figure 2a**). At the patient level, we observed high concordance among patients with similar tumor types, except in a few cases where samples had insufficient malignant cells (**Figure 2b**). However, basic analyses revealed palpable differences but were insufficient due to the high inter-patient and inter-tumor heterogeneity of malignant cells^15^. Therefore, we employed a more sophisticated method: non-negative matrix factorization (NMF)-based metaprogram (MP) generation, adopted from previous studies^15,16^.

**Fig. 2.**
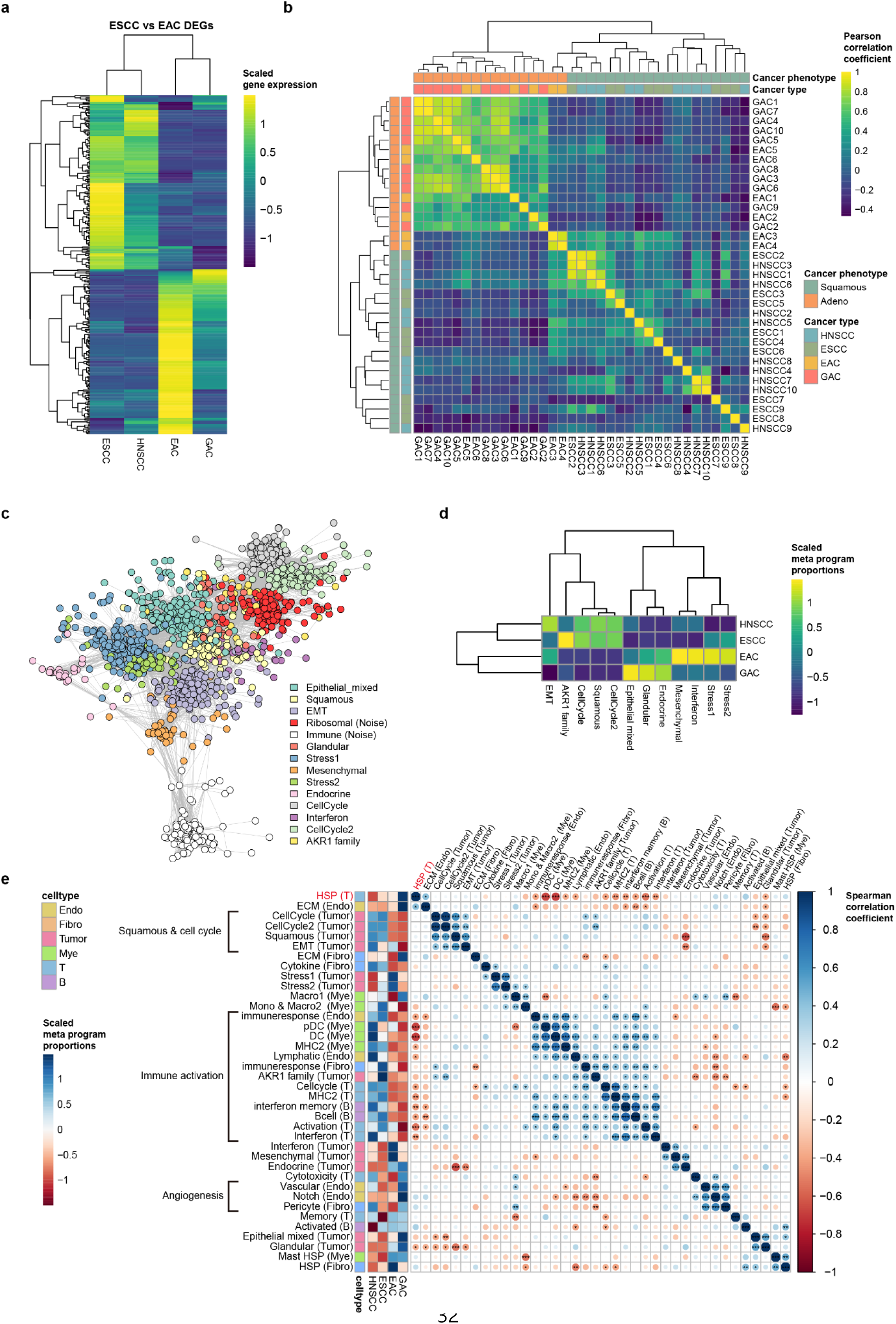
Metaprogram analysis. **a**, Heatmap of DEG enrichment from comparing malignant epithelial cells from ESCC and EAC. Rows and columns are clustered with hierarchical clustering. Colors indicate z-scaled averaged normalized expression values for each tumor type. **b**, Heatmap of patient similarities. The colors indicate the Pearson correlation coefficient. Rows and columns are indicated by histological phenotypes and cancer types. **c**, The Co-occurrence network with genes from NMF modules represented in the Fruchterman-Reingold layout. Each node indicates a gene, and the nodes are colored by their metaprograms. **d**, Heatmap for proportions of ‘on’ cells for each meta program calculated for each tumor type. Proportions are z-scaled column-wise. The rows and columns are clustered with hierarchical clustering. **e**, Heatmap for spearman correlations among patients with their proportions of ‘on’ cells for each meta program. The color indicates spearman correlation coefficient (\**P* < 0.05; \*\**P* < 0.01; \*\*\**P* < 0.001). Each row is additionally indicated for tumor type-wise ‘on’ cell proportions and cell types.

Initially, 14 malignant MPs were generated by clustering genes from the NMF modules based on their co-occurrence (**Figure 2c**, **Supplementary Table 4**). Each MP was annotated through manual curation, pathway enrichment analysis, and comparison with previously generated MPs (**Supplementary Table 5**, **Supplementary Figure 2a**). We then calculated proportion of malignant cells that were ‘on’ for each MP. Based on the per-patient proportions of cells active for each MP, we measured overall enrichment of each MP for each cancer type (**Figure 2d**). As expected, we observed separations of cancer types based on their histological origins, represented by the Squamous and Glandular MPs. Patient-level enrichment showed similar overall results (**Supplementary Figure 2b**), indicating clear distinctions of malignant epithelial cells between cancer types with different histological origins.

We also noted enrichment of Cell cycle MPs for HNSCC and ESCC (**Figure 2d**). MPs with cell cycle-related genes are often enriched across all tumor types^15^. However, the specific enrichment of Cell cycle MPs might reflect higher frequencies of cyclin-dependent kinase alterations in HNSCC and ESCC compared to adenocarcinoma^17^. For EAC and GAC, we observed enrichment of Endocrine MP, reflecting the formation of glandular architecture in tumor cells for these cancers^18^. For ESCC and partially for HNSCC, an MP enriched with various AKR1 genes (*AKR1C3*, *AKR1C1, AKR1B10, AKR1C2*) was specifically observed. AKR1 family genes are known to activate the PI3K/AKT signaling pathways^19^, which are highly activated in these cancers^20,21^. Overall, MP analysis revealed that while histology-based meta programs remain the most dominant factors, additional MPs explain the distinct states of cancer cells between ESCC and EAC.

### Distinction of TME between ESCC and EAC by immune activity

To analyze the possible connections between cancer cells and TME compartments, we further dissected immune and stromal cells in TME by generating MPs from each major cell type; T cells, myeloid cells, B cells, fibroblasts, and endothelial cells (**Supplementary Figure 2c**). We also measured proportions of ‘on’ cells for each major cell type with its matching MPs and calculated pair-wise correlations among proportions of all MPs (**Figure 2e**). Tumor MPs related to histology of epithelial cells, such as the Squamous and Glandular MPs, showed negative correlations, reflecting their opposite enrichment based on cancer types. Additionally, we observed a large cluster of immune-activating MPs, including Fibroblast- and endothelial-immune response MPs, Myeloid-DC MPs and T-activation and T-interferon MPs. Among tumor MPs, the AKR1 family gene MP was included in the immune-activating cluster, indicating possible importance of these genes in the TME. The immune-activating MP cluster suggested that regulation of immune activation in the TME is coordinated across different cell types. Interestingly, heat-shock protein (HSP) MP from T cells showed a negative correlation with immune-activating MPs. The MPs in the immune-activating cluster were mostly enriched for squamous cell carcinomas, while the anti-correlated T-HSP MP (T cells with high HSP signals) was enriched for GAC. High HSP signals for T cells were recently identified as stress-response T cell signatures, which are known to possibly turn the TME into a cold tumor, negatively affecting the patient’s response to immunotherapy^22^. GAC showed high enrichment of various ECM-building and angiogenesis related fibroblast and endothelial cell MPs, indicating pro-tumor activities from those cells^23^. These findings suggest that immune-activating factors in the TME are more enriched in both HNSCC and ESCC, while pro-tumoral stromal cell activities prevail for GAC. EAC samples display intermediate patterns of MP enrichment, potentially reflecting partial characteristics of ESCC and GAC.

### ESCC and EAC differ in the ratio of *CXCL13*^+^ and stress-responsive CD8^+^ T cells

To further understand the TME, we conducted in-depth analyses of each major cell type using a subclustering approach. From the integrated immune and stromal cell population, we isolated CD8^+^ T cells. Using unbiased clustering and marker-based cell type annotation, we identified six sub-subs of CD8^+^ T cells (**Figure 3a**). We annotated two memory CD8^+^ T cell subsets without effector functions as the naïve & memory subset and the *GZMK*^+^ memory subset, based on *GZMK* expression. Three subsets with effector functions were identified and named as IFN high, Effector, and Exhausted CD8^+^T cell subsets, characterized by the expression of interferon-related, cytotoxicity-related, and exhaustion-related genes, respectively (**Figure 3b, Supplementary Table 7**). We scored each cell with an exhaustion signature (**Supplementary Table 5-6**) and observed increasing scores for both signatures from naïve to exhausted CD8^+^ T cell subsets (**Supplementary Figure 3a**). The cell type composition analysis showed that HNSCC and ESCC had more CD8^+^ T cells with effector functions, while EAC and GAC samples had significantly more HSP-high CD8^+^ T cells (**Figure 3c**).

**Fig. 3.**
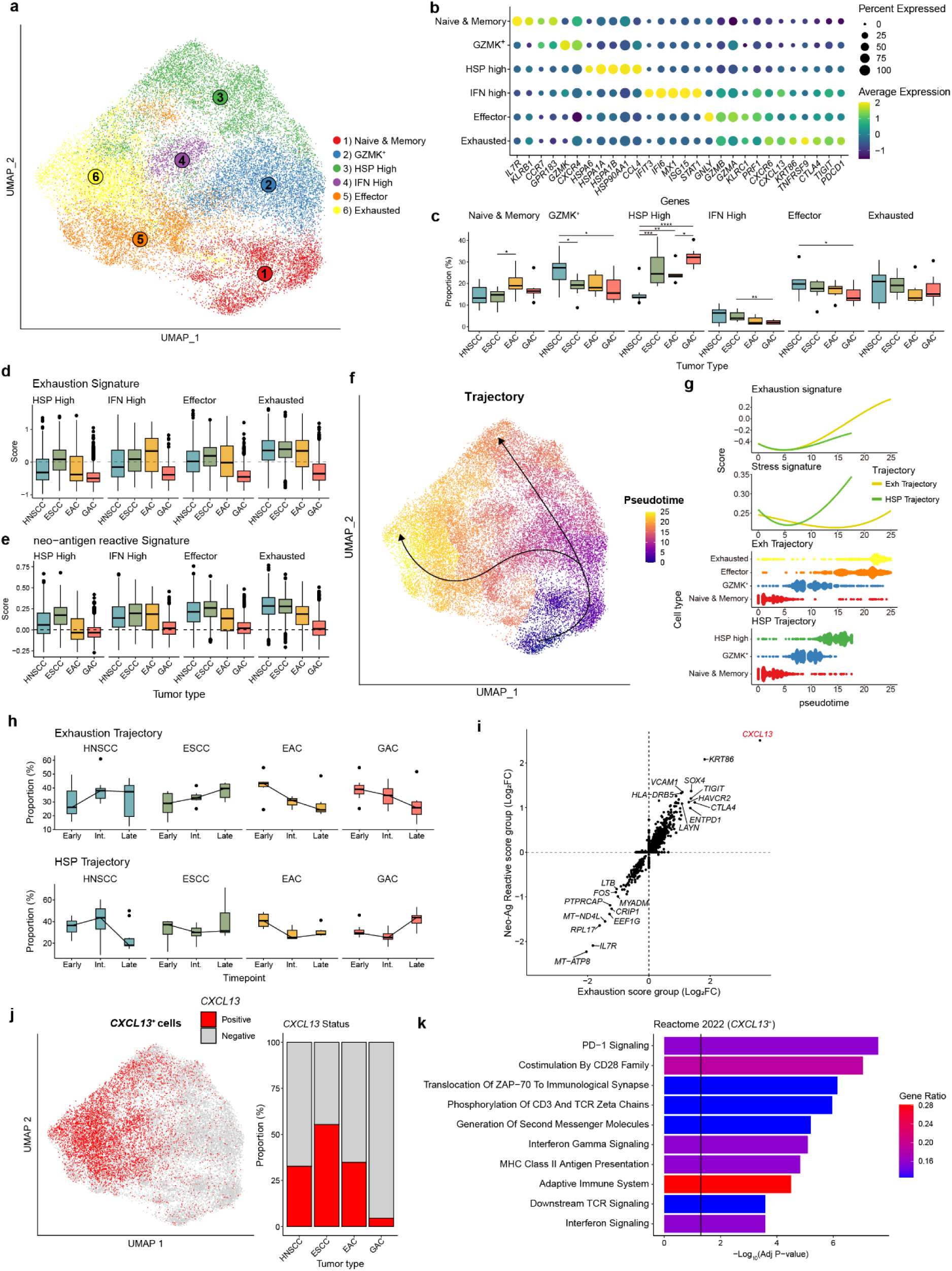
Distinct subsets of CD8^+^ T cells. **a,** UMAP of CD8^+^ T cell subsets from subclustering analysis. HSP; heat-shock protein, IFN; interferon. **b**, Dotplot of markers genes for each subset of CD8^+^ T cells. The color indicates scaled average expression for each subset and the sizes indicate percentage of cells that are expressing each gene. **c**, Proportion of each subset for each patient. The proportions are visualized by the boxplots for each cancer type and each CD8^+^ T cell subset. **d-e**, Boxplots of (d) exhaustion scores and (e) neo-antigen reactive scores from each cell. The boxplots are visualized for each subset and each cancer type. **f**, UMAP of CD8^+^ T cells with pseudotimes and their trajectories into exhausted CD8^+^ T cells or HSP-high CD8^+^ T cells. The colors indicate pseudotimes for each cell. **g**, (Top) Exhaustion signature scores and stress response signature scores along the exhaustion and HSP trajectories. (Bottom) Pseudotimes of cells along exhaustion and HSP trajectories colored by their cell types. **h**, Boxplots of proportions of cells for each cancer type in early, intermediate, and late stages of each trajectory for each patient. **i**, DEGs annotated with fold changes calculated between cells with high and low exhaustion signature scores (x-axis) and between cells with high and low neo-antigen reactive signature scores (y-axis). **j**, (Left) UMAP of CD8^+^ T cells with *CXCL13* expression. (Right) Proportions of CD8^+^ T cells for each tumor type with or without *CXCL13* expression (colored in red or grey, respectively). **k**, Pathway analysis using Reactome database with DEGs from comparing *CXCL13*^+^ CD8^+^ T cells to *CXCL13*^−^ CD8^+^ T cells. Colors of the bar indicate ratio of genes from the DEGs that are in gene list for each pathway term. Black vertical lines indicate q-value threshold of 0.05.

To exclude possibility of different batches affecting the clustering results, we utilized the T cell-specific reference-based annotation method, projecTIL^24^. With its default CD8^+^ T cell reference, we annotated CD8^+^ T cells in each tumor type separately. The results from projecTIL showed a higher proportion of exhausted T cell populations for ESCC, while the opposite trend was observed for central memory T cells (**Supplementary Figure 3b**). This suggests that degree of T cell exhaustion is an important factor distinguishing these tumor types. Indeed, both adenocarcinomas tended to show lower exhaustion signal even within the same sub-population (**Figure 3d**). To determine whether these exhausted populations were highly tumor-reactive, we scored cells using tumor neo-antigen reactive signatures (**Supplementary Table 5-6**). Similar to the exhaustion signal, squamous carcinoma types exhibited the highest tumor reactive scores **(Figure 3e)**. Furthermore, the scores from these two signatures were significantly correlated (**Supplementary Figure 3c**).

We noticed possible existence of more bystander CD8^+^ T cells for EAC and GAC indicated by having fewer activated populations despite having more CD8^+^ T cells. In addition to the exhausted T cells, another noticeable population was T cells with high HSP. A recent single-cell pan-cancer study identified these HSP-high T cells as stress-responsive populations, which could be poor indicators of ICB responses^22^. This suggests that the tumor-reactive exhausted T cells and stress-responsive T cells might have functionally opposite roles. Therefore, we conducted trajectory analysis on CD8^+^ T cells for further evaluation of these populations. Using monocle3^25^, we found that these populations formed separate branches of developmental trajectories: the exhaustion trajectory and the HSP trajectory. (**Figure 3f**, **Supplementary Figure 3d-e**). We confirmed that these cells align well along the pseudotime following expected developmental and activation paths (**Supplementary Figure 3f**). Using exhaustion signature and stress-related signature (**Supplementary Table 5-6**), we showed that exhaustion signature increases along the exhaustion trajectory while stress-related signature increases along the HSP trajectory (**Figure 3g, top**). Along the pseudotime of each trajectory, we identified corresponding cell populations to align well with increasing pseudotime of both trajectories (**Figure 3g, bottom**). To identify how proportions of cells for each cancer type changes along the trajectory, we calculated proportion of cells in early to late stages of the trajectory. For exhaustion trajectory, there were increasingly more populations at the late stage for HNSCC and ESCC but opposite for EAC and GAC (**Figure 3h, top**). In contrast, the HSP trajectory showed increasing population for GAC while sharp decrease between intermediate to late stages for HNSCC and small decrease from the early to late stages for ESCC. Although EAC showed no decrease from the intermediate to late stage, a larger population at the early stage of both exhaustion and HSP trajectories resulted in decreased populations of EAC for both trajectories (**Figure 3h, bottom**). Overall, trajectory analysis also shows strong trend of increasing exhaustion for HNSCC and ESCC, a sharp increase in HSP populations for GAC, and EAC cells displaying intermediate characteristics between squamous cancer types and GAC.

The observed differences in CD8^+^ T cell populations may have a possible connection to ICB responses. The infiltration and activation status of T cells, along with their tumor reactivity, are crucial in the formation of ‘hot tumors’ or ‘cold tumors’, which are known to be associated with ICB efficacy^26^. Given the varying degree of exhaustion and tumor-reactivity within the exhausted T cell populations, we sought to identify the genes contributing to this variation. We performed DEG analysis between cells with high and low exhaustion scores and neoantigen reactive scores (**Supplementary Table 8**). Among the genes related to both T cell exhaustion and tumor reactivity, *CXCL13* ranked the highest in both comparisons (**Figure 3i**). We confirmed that *CXCL13* expression is highly enriched in effector and exhausted T cell populations, with GAC samples showing a very low percentage of cells expressing *CXCL13* (**Figure 3j**, **Supplementary Figure 3g**). Effector and exhausted CD8^+^ T cells without *CXCL13* expression showed low signature scores for exhaustion and neo-antigen reactivity, comparable to other non-effector CD8^+^ T cells (**Supplementary Figure 3h**). Furthermore, pathway analysis for *CXCL13*^+^ T cells indicated high levels of T cell infiltration and activation, with pathways related to IFN-γ signaling, T cell activation, and antigen processing (**Figure 3k, Supplementary Figure 3i**). Overall, CD8^+^ T cell analysis identified major differences among four cancer types, particularly *CXCL13*^+^ T cells and stress-responsive T cells. These differences may be connected to immune mechanisms that differentiate ICB responses among the four cancer types.

### ESCC and EAC may differ in capability of TLS formation

Along with CD8^+^ T cells, CD4^+^ T cells are also involved in various immunotherapy-related immune activities, including helping or regulating CD8^+^ T cells and B cells, and exerting direct cytotoxicity on tumor cells^27^. In our datasets, the major immune mechanism related to ICB response for CD4^+^ T cells was the formation of TLS with follicular helper T (Tfh) cells. Overall, we identified various CD4^+^ T cell subsets with subclustering analysis (**Figure 4a**). The subsets include naïve and memory populations with *CCR7* and *IL7R* expression, a cytotoxic subset (CTL) with *GZMA* and GZMB expression, Tfh with *CXCL13* and *TOX*, HSP-high subsets, and several subsets of regulatory T cells (**Supplementary Figure 4a, Supplementary Table 9**). From these subsets, we further analyzed Tfh cells, which are important components of TLS alongside germinal center B cells (GCBs), both considered critical for ICB responses^28^. ESCC showed the highest proportions of Tfh cells (**Figure 4b, Supplementary Figure 4b-c**) and Tfh cells from HNSCC and ESCC expressed more *CXCL13* (**Figure 4b**). This finding aligns with the recent reports highlighting the existence and importance of TLS formation in ESCC^29,30^. Furthermore, CXCL13-expressing Tfh cells are known to induce the maturation of early TLS into follicle-formed TLS through B cell recruitment and maturation^31^.

**Fig. 4.**
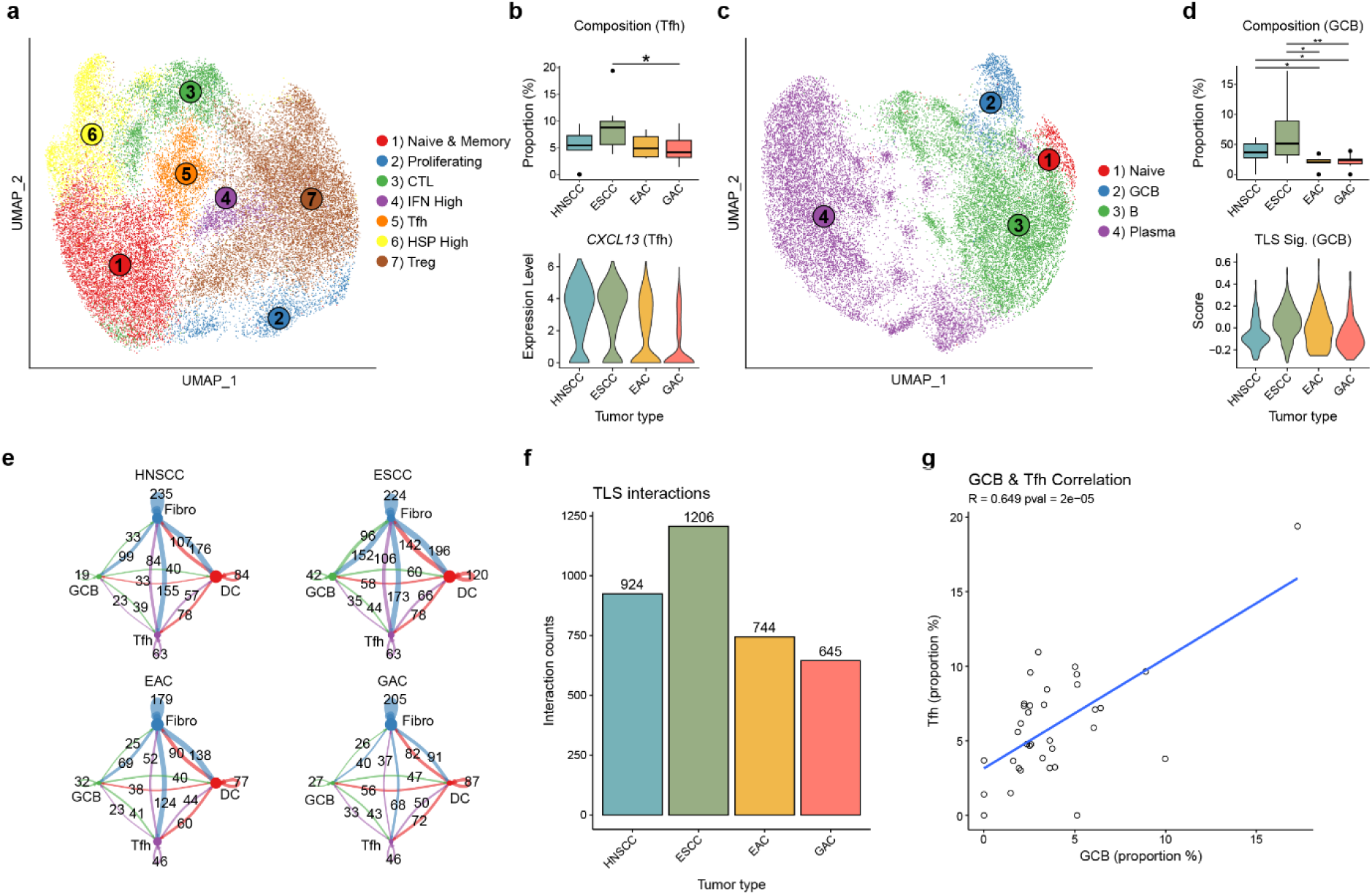
Subpopulations of CD4^+^ T cells and B cells. **a,** UMAP of CD4^+^ T cells with annotated subpopulations. CTL; cytotoxic lymphocyte, IFN; interferon, Tfh; follicular helper T cell, HSP; heat-shock protein, Treg; regulatory T cell. **b**, (Top) Proportion of Tfh out of CD4^+^ T cells for each patient. The proportions are visualized by the boxplots for each tumor type. (Bottom) Violin plots for *CXCL13* expression from Tfh. **c**, UMAP of B cells with annotated subpopulations. GCB; germinal center B cell. **d**, (Top) Proportion of GCBs out of B cells for each patient. The proportions are visualized by the boxplots for each tumor type. (Bottom) Violin plots for TLS signature scores from GCB. **e**, Number of interactions among fibroblasts (blue), GCBs (green), Tfh (purple), and DCs (red). The colors of lines indicate that the interactions are counted with the matched cell types as the senders. Fibro; fibroblasts, DC; dendritic cells. **f**, Total interaction counts for each tumor type among the cell types in Fig. 4e. Self-interaction counts were excluded. **g**, Correlation between proportions of GCB (of B cells) and Tfh (of CD4^+^ T cells) for each patient.

From subclusters of B cells and plasma cells (**Figure 4c**), we identified GCBs characterized by the expression of *RGS13* and *ACTB* (**Supplementary Figure 4d, Supplementary Table 10**). As expected, GCBs showed the highest expression of TLS signature (**Supplementary Figure 4e, Supplementary Table 5-6**). Among the cancer types, ESCC exhibited the highest proportions of GCBs with higher TLS signatures (**Figure 4d**). We also calculated cell-cell interactions among compartments of TLS including DCs, GCBs, fibroblasts, and Tfh cells (**Figure 4e**). Among these cell types, we observed stronger interactions in HNSCC and ESCC (**Figure 4f**). Across all samples, we observed a positive correlation between the proportions of Tfh cells and GCBs (**Figure 4g**). Due to higher *CXCL13* expression by Tfh cells and enrichment of GCBs, both HNSCC and ESCC appeared to indicate the formation of TLS with these populations.

### ESCC and EAC differ in the ratio of *CXCL9*^+^*CXCL10*^+^ and *MARCO*^+^ TAMs

Myeloid cells are also highly connected to ICB responses through their role in regulating T cells as antigen presenting cells (APCs), such as dendritic cells (DCs) and macrophages. Furthermore, TAMs interact with T cells via their expression of myeloid immune checkpoint ligands^32^. We performed subclustering analysis of myeloid cells and identified monocytes, myeloid-derived suppressor cells (MDSCs), TAMs, three subtypes of DCs, and mast cells (**Figure 5a**). Monocytes and MDSCs share monocyte-related markers such as *CD14* and *S100A8,* but MDSCs express chemokines and interferon-related genes. DCs modulate T cell activation and suppression and are highly implicated in ICB response^33^. We identified three subsets of DCs, conventional DCs (cDCs) with *CD1C* and *CLEC10A* expression, more mature and activated DCs (Act. DCs) with DC maturation markers such as *CCR7* and *FSCN1*, and plasmacytoid DCs (pDCs) with *GZMB* and *JCHAIN* expression (**Figure 5b, Supplementary Table 11**). Cell type abundance analysis showed fewer mature myeloid cells in EAC, while HNSCC and ESCC tended to have more cDC and activated DC subsets (**Figure 5c**). Pathway analysis of the activated DC marker genes revealed various pathways related to positive regulation of T cells and cytokine productions (**Supplementary Figure 5a**). Furthermore, the proportions of activated DCs positively correlated with TME MPs related to T cell activation and interferon production and negatively correlated with the glandular tumor MP and the heat-shock response T cell MP, which we speculated to be a negative indicator of ICB responses (**Figure 5d**). *BATF3* expression by the activated DCs, necessary for effector T cell activation^34^, was almost absent in GAC (**Supplementary Figure 5b**). Along with decreased proportions of activated DCs in EAC and GAC, the absence of *BATF3* expression could be connected to the low activation status of T cells for these cancer types.

**Fig. 5.**
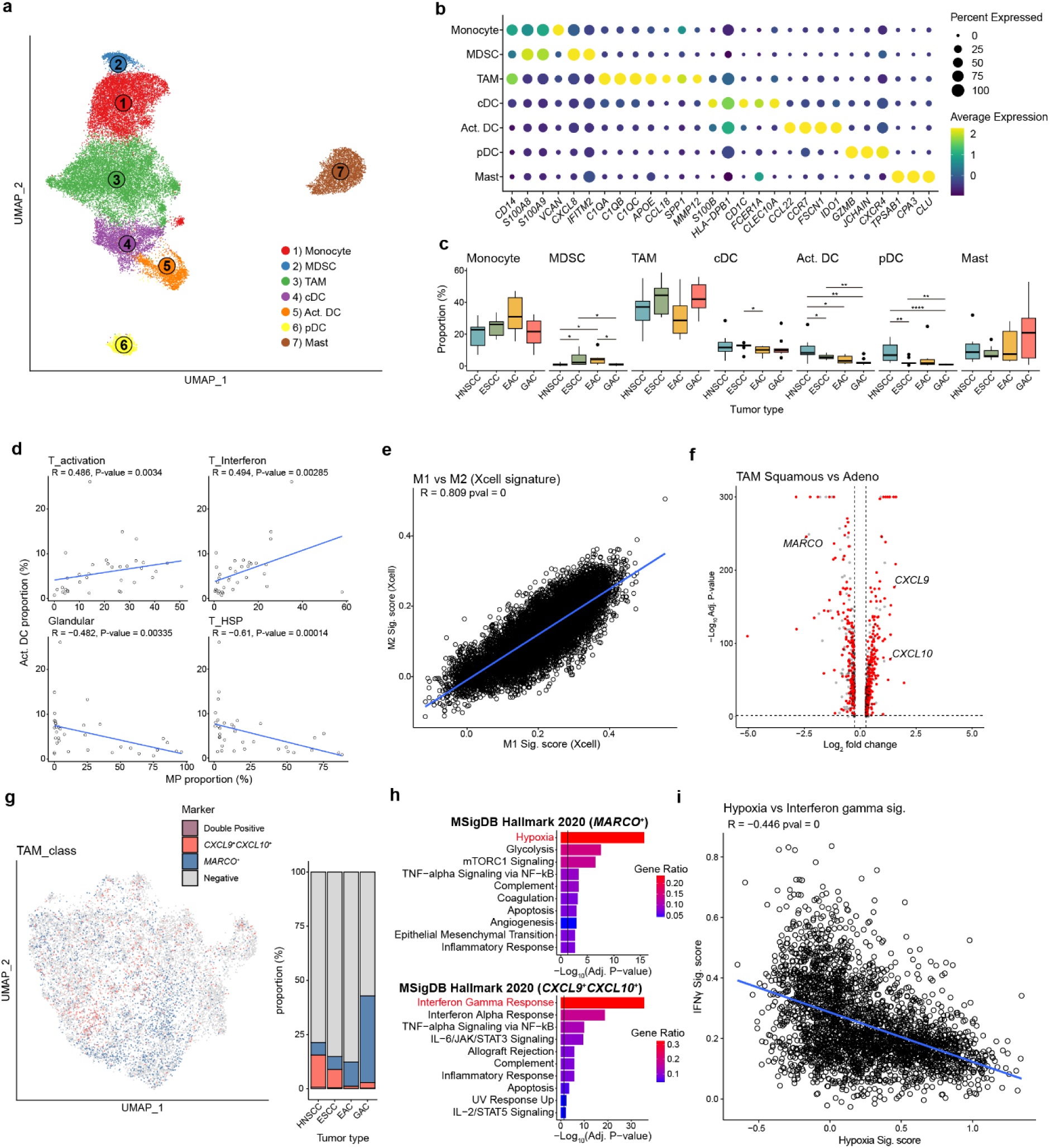
Exclusive TAM subsets and their opposite functions. **a,** UMAP plot of myeloid cells with annotated subpopulations. MDSC; myeloid-derived suppressive cells, TAM; tumor-associated macrophages, cDC; conventional DCs, Act. DC; activated DCs, pDC; plasmacytoid DCs. **b**, Dotplot of marker genes for each subpopulation of myeloid cells. The color indicates scaled average expression for each subpopulation and the sizes indicate percentage of cells that are expressing each gene. **c**, Proportion of each subpopulation per patient, visualized by boxplots for each cancer type and myeloid cell subpopulation. **d**, Correlation between cellular proportions of four metaprograms (T_activation, T_interferon, Tumor_glandular, T_HSP) and proportions of activated DCs (of myeloid cells). **e**, Correlation between each cell’s M1 and M2 macrophage polarization signature scores. **f**, DEGs between squamous cell carcinomas (ESCC and HNSCC) and adenocarcinomas (EAC and GAC). Genes colored red are DEGs for both ESCC vs EAC and squamous vs adenocarcinomas. **g**, UMAP plot of TAMs that are colored by expression of *CXCL9*/*CXCL10* and *MARCO*. **h**, Pathway enrichment analysis with MsigDB hallmark with DEGs from (Top) *MARCO*^+^ TAMs and (Bottom) *CXCL9*^+^*CXCL10*^+^ TAMs. Colors of the bar indicate ratio of genes from the DEGs that are in gene list for each pathway term. Black vertical lines indicate q-value threshold of 0.05. **i**, Correlation between scores from hypoxia signature and IFN-γ signature for TAMs. TAMs not expressing *MARCO*, *CXCL9*, and *CXCL10* are filtered.

For more detailed annotation of TAMs, we initially attempted to identify polarization status of each TAM cluster. However, M1 and M2 signatures were not clearly distinguishable among clusters and even showed positive correlations among all TAMs (**Figure 5e**). Instead of canonical polarization markers, there could be distinct populations of TAMs depending on epithelial origins of each cancer type. We compared TAMs from HNSCC and ESCC samples to those from EAC and GAC samples. Among DEGs between these two groups, we identified *MARCO* to be enriched for EAC and GAC, while *CXCL9* and *CXLC10* were enriched in HNSCC and ESCC (**Figure 5f**, **Supplementary Figure 5c, Supplementary Table 12**). These genes might be important markers distinguishing TAM sub-populations since they were highly exclusive by having only 0.4% of TAMs with both markers (**Figure 5g**). As expected, HNSCC and ESCC samples had more *CXCL9*^+^*CXCL10*^+^ TAMs, while EAC and GAC TAMs were mostly *MARCO*^+^ TAMs. We then identified the functions of each TAM subset. Pathway analysis indicated that *CXCL9*^+^*CXCL10*^+^ TAMs were enriched with IFN-γ pathways, a positive indicator of ICB response, while *MARCO*^+^ TAMs were enriched with hypoxia pathways (**Figure 5h**). Recent studies associate hypoxia with resistance to ICB response by interfering with other immune populations^35^. Furthermore, a recent study on HNSCC also identified two clinically important TAM populations: *CXCL9*^+^ TAMs and *SPP1*^+^ TAMs^36^. *SPP1* is one of the markers of *MARCO*^+^ TAMs. Therefore, we concluded that these TAMs have opposite functions related to ICB responses. We also scored each TAM with IFN-γ-related genes as a positive signature for ICB and hypoxia-related genes as a negative signature. These signatures showed similar enrichment patterns as the pathway analysis (**Supplementary Figure 5d**) and had negative correlations, as expected from the exclusiveness of the markers (**Figure 5i**).

### Immune signatures distinguishing esophageal cancer subtypes predict ICB responses

Two cell types with the most noticeable patterns among the four cancer types were CD8^+^ T cells and TAMs. We focused on the *CXCL13*^+^CD8^+^ T cells, HSP-high CD8^+^ T cells, *CXCL9*^+^*CXCL10*^+^ TAMs, and *MARCO*^+^ TAMs due to their possible implications in ICB responses. A recent study have shown that CXCL13^+^LAG3^+^ T cells localize in CXCL9 and CXCL10 high patches, leading to the formation of hot tumor^37^. Moreover, IFN-γ-dependent productions of CXCL9 and CXCL10 by macrophages is necessary for the therapeutic efficacy of checkpoint blockade^38^. Conversely, MARCO is a myeloid checkpoint that promotes tumors, and its involvement in immunotherapy efficacy is related to hypoxia^32,39^. HSP-high T cells have recently been implicated as negative indicators of immunotherapy response with their localization in hypoxia-rich regions^22^.

Since *CXCL13*^+^CD8^+^ T cells and *CXCL9*^+^*CXCL10*^+^ TAMs were both enriched for HNSCC and ESCC, and both subtypes shared IFN-γ-related pathways, we performed cell-cell interaction analysis between these two subtypes compared to *CXCL13*^−^CD8^+^ T cells and *CXCL9*^−^*CXCL10*^−^ TAMs. Using CellChat^40^ for cell-cell interaction inference, we found more interactions between *CXCL13*^+^CD8^+^ T cells and *CXCL9*^+^*CXCL10*^+^ TAMs (**Figure 6a**). To confirm the coordinated functions and localization of these populations, we quantified co-abundance of these two subtypes. HNSCC and ESCC patients both showed significant co-abundance of these subtypes while EAC and GAC patients did not (**Figure 6b**). Similarly, we analyzed cell-cell interactions and co-abundance of HSP-high T cells with *MARCO*^+^ TAMs, as both subtypes are related to hypoxia in immune cells and are negative indicator of ICB responses. We observed more interactions between HSP-high CD8^+^ T cells and *MARCO*^+^ TAMs compared to *MARCO*^−^ TAMs (**Figure 6c**). GAC patients showed significant co-abundance of these two subtypes (**Figure 6d**).

**Fig. 6.**
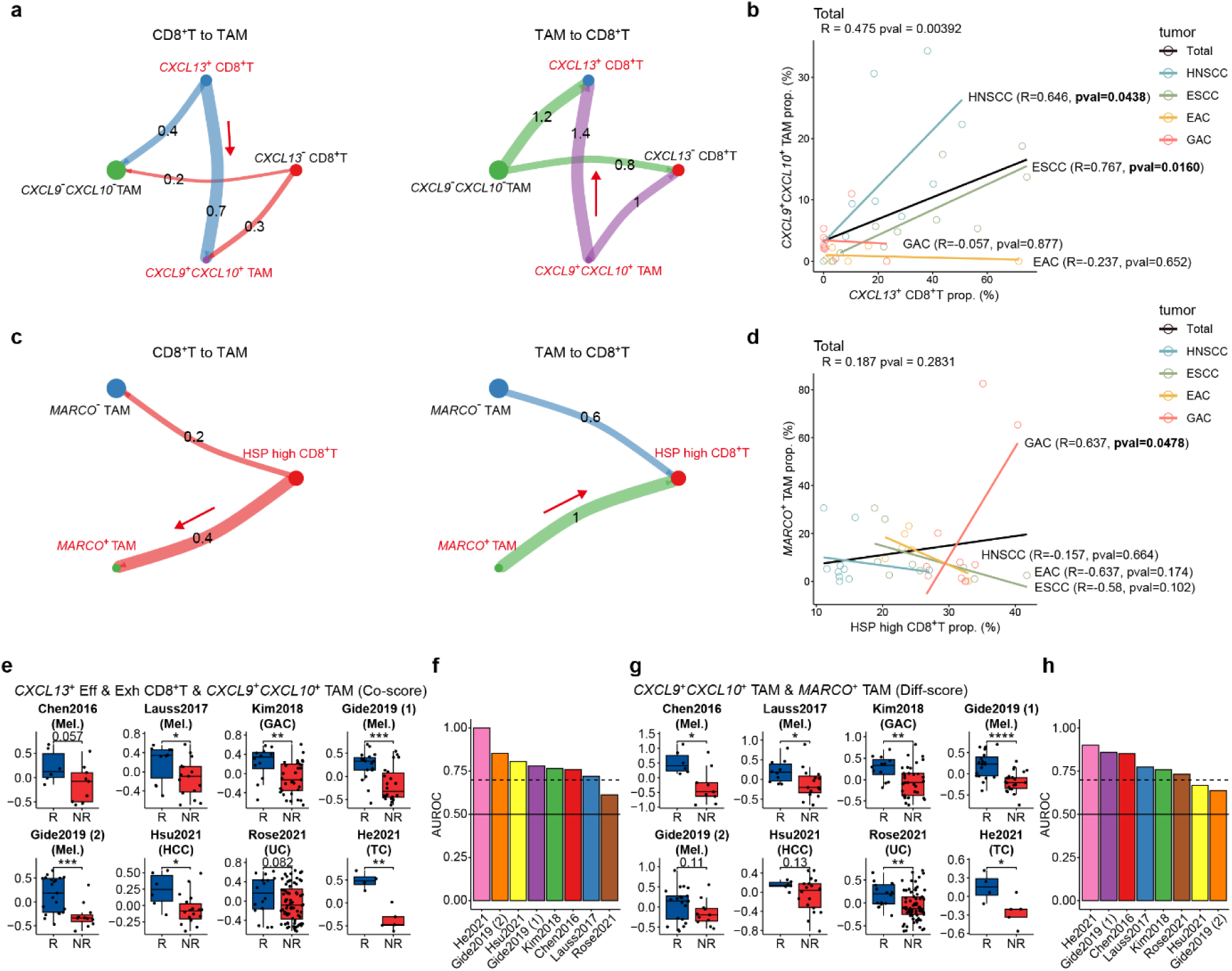
CD8+T cell and TAM subset signatures predictive for ICB responses. **a,** Circle plots of Intercellular communication probability generated by Cellchat among *CXCL13*^+^CD8^+^ T cells, *CXCL13*^−^CD8^+^ T cells, *CXCL9*^+^*CXCL10*^+^ TAMs, and *CXCL9*^−^*CXCL10*^−^ TAMs. The arrows indicate interactions between *CXCL13*^+^CD8^+^ T cells and *CXCL9*^+^*CXCL10*^+^ TAMs. We visualized only interactions from (Left) CD8^+^ T cells to TAMs or (Right) TAMs to CD8^+^ T cells. **b**, Correlation between proportion of *CXCL13*^+^CD8^+^ T cells and *CXCL9*^+^*CXCL10*^+^ TAMs. **c**, Circle plots of intercellular communication probability among HSP-high CD8^+^ T cells, *MARCO*^+^ TAMs, and *MARCO*^−^ TAMs. The arrows indicate interactions between HSP-high CD8^+^ T cells and *MARCO*^+^ TAMs. We visualized only interactions from (Left) CD8^+^ T cells to TAMs and (Right) TAMs to CD8^+^ T cells. **d**, Correlation between proportion of HSP-high CD8^+^ T cells and *MARCO*^+^ TAMs. **e**, Boxplots of GSVA scores calculated by averaging scores with gene signatures from *CXCL13*^+^ effector and exhausted CD8^+^ T cells and *CXCL9*^+^*CXCL10*^+^ TAMs (Co-score). The scores were calculated for each patient and grouped by ICB response (R, responder; NR, non-responder). Mel., melanoma; HCC, hepatocellular carcinoma; UC, urothelial carcinoma; TC, thymic carcinoma. **f**, AUROC scores for prediction of ICB responders with Co-score. **g**, Boxplots of GSVA scores calculated by subtracting a score with gene signatures from *MARCO*^+^ TAMs from a score with *CXCL9*^+^*CXCL10*^+^ TAMs (Diff-score). The scores were calculated for each patient and grouped by ICB response. **h**, AUROC scores for prediction of ICB responders with Diff-score.

To validate the involvement of these cell subsets in immunotherapy responses, we used bulk RNA-seq datasets from eight cohorts across diverse cancer types, including melanoma, gastric cancer, urothelial cancer, thymic cancer, liver cancer (**Supplementary Table 13**). These datasets included patient information on ICB responses and samples collected prior to treatment. We generated cellular signatures based on DEGs by comparing *CXCL13*^+^ effector and exhausted CD8^+^ T cells, *CXCL9*^+^*CXCL10*^+^ TAMs, and *MARCO*^+^ TAMs to their negative counterparts (**Supplementary Table 14**). We excluded HSP-high CD8^+^ T cells as their DEGs were mostly genes for HSPs, which are expressed by a wide range of cell types, making them indistinguishable in bulk transcriptomes. We calculated co-enrichment scores (co-score) by averaging enrichment scores of two cellular signatures. We also determined differences between two cellular signature scores as difference score (diff-score). For co-score, we chose the signatures for *CXCL13*^+^CD8^+^ T cells and *CXCL9*^+^*CXCL10*^+^ TAMs due to their higher interactions. The co-scores from these two cellular signatures showed significant differences between responders and non-responders in six out of the eight cohorts (**Figure 6e**) and were highly predictive for responders (AUROC > 0.7) in seven out of the eight cohorts (**Figure 6f**). For the diff-score, we focused on differences between the two TAM subset signatures: *CXCL9*^+^*CXCL10*^+^ TAMs and *MARCO*^+^ TAMs. The diff-scores also showed significant differences between responders and non-responders for six out of the eight cohorts (**Figure 6g**) and were predictive for responders in six out of eight cohorts (**Figure 6h**). These results suggest that cellular signatures distinguishing esophageal cancer subtypes can predict ICB response across diverse cancer types.

## Discussion

In the present study, we compared two subtypes of esophageal cancer, ESCC and EAC, at single-cell resolution. To enhance the distinctness between the subtypes, we supplemented additional cancer types near the esophagus known to be molecularly closer to each subtype: HNSCC for ESCC and GAC for EAC. Through the comparative single-cell analysis of the four cancer types, we confirmed distinctions between the two subtypes of esophageal cancer and their shared similarities with nearby cancer types in both cancer cells and the TME.

Our results show that malignant epithelial cells have the most distinct separations based on their histological origins, followed by varying cancer cell states for each tumor type. For immune cells, we found that hot tumor-related immune subsets such as *CXCL13*^+^CD8^+^ T cells and *CXCL9*^+^*CXCL10*^+^ TAMs, which share pathways of IFN-γ responses, are enriched in HNSCC and ESCC. Conversely, cold tumor-related immune subsets such as HSP-high CD8^+^ T cells and *MARCO*^+^ TAMs, which are related to hypoxic environments, are more enriched in EAC and GAC. These associations between immune subsets and cancer types near the esophagus may explain the higher response rate to immunotherapy in HNSCC and ESCC compared to EAC and GAC. Finally, we found that expression signatures derived from these IFN-γ- and hypoxia-related immune cell subsets are predictive for ICB responses in diverse cancer types.

Recapitulating bulk tissue-level molecular distinctions between ESCC and EAC and their respective similarities to HNSCC and GAC, malignant cells from HNSCC and ESCC resembled squamous epithelial cells, while malignant cells from EAC and GAC were enriched with glandular epithelial cells. To further discover various cancer cell states, we used NMF analysis to generate MPs that could represent heterogeneous malignant functions. Along with histology-related MPs, we annotated several functional MPs, such as those related to cell cycle, that could reflect the effects of malignant cells on TME. Thus, the single-cell models of cancer cells, accounting for inter-tumor heterogeneity, align well with tissue-level models of tumors for the same cancer types.

Unlike malignant cells, we expected less superficial differences in immune cells but rather subtle differences in underlying immune mechanisms. To capture these differences, we conducted subclustering analyses of major immune cell types. For CD8^+^ T cells, we identified two diverging developmental pathways for exhausted CD8^+^ T cells and HSP-high CD8^+^ T cells. Along with opposing trajectories, exhausted yet neo-antigen reactive CD8+ T cells and stress-responsive HSP CD8^+^ T cells indicated possible opposite functions during immunotherapy. We identified that populations of exhausted CD8^+^ T cells with *CXCL13* expression would have more neo-antigen reactivity and enrichment of IFN-γ signatures. Among four cancer types, ESCC followed by HNSCC showed stronger enrichment of *CXCL13*^+^ CD8^+^ T cells with more transition into exhaustion trajectory. Conversely, we identified transition of naïve GAC CD8^+^ T cells to HSP-high CD8^+^ T cells. However, EAC samples exhibited characteristics of both squamous carcinomas and GAC. We also noticed differential enrichment of TAM subsets among the four cancer types. We identified *CXCL9*^+^*CXCL10*^+^ TAMs and *MARCO*^+^ TAMs by comparing TAMs from HNSCC and ESCC to those from EAC and GAC. Based on their enriched pathways and literature evidence, we concluded possible opposite roles for ICB responses from IFN-γ-related *CXCL9*^+^*CXCL10*^+^ TAMs and hypoxia-related *MARCO*^+^ TAMs. Furthermore, we analyzed cell-cell interactions among these matching populations. The results suggested both higher interactions between *CXCL13*^+^ CD8^+^ T cells and *CXCL9*^+^*CXCL10*^+^ TAMs and between HSP-high CD8^+^ T cells and *MARCO*^+^ TAMs.

This study has some limitations. Due to the rare incidence of EAC, the study has a limited sample sizes for this subtype. However, the analytical strategy of including nearby cancers for comparison partially overcome this limitation by making the differences between ESCC and EAC more pronounced and identifiable. Additionally, due to lack of ICB-treated patients, our results may not directly indicate involvement in ICB responses. To address this limitation, we validated the cellular subsets distinguishable between subtypes with varying ICB responses using public transcriptome data with ICB-treated cohorts for diverse cancer types.

## Materials and Methods

### Tumor specimen collection and sample preparation

Patient samples were collected from individuals diagnosed with esophageal cancer (ECC) who underwent surgical resection at Severance Hospital in Seoul, Republic of Korea, between 2019 and 2021. Tumors were cut into small pieces and processed using a gentleMACS dissociator (Miltenyi Biotec, Gladbach Bergisch, Germany, Cat#130-093-235) and enzymatically digested using the Human Tumor Dissociation Kit (Miltenyi Biotec, Cat#130-095-929) following the manufacturer’s protocols. After incubating for 1 hour at 37°C, samples were filtered through a 70-µm MACS SmartStrainer (Miltenyi Biotec, Cat#130-098-462) into RPMI-1640 medium (Corning, Inc., Corning, NY, USA) supplemented with 10% fetal bovine serum (Biowest, Riverside, MO, USA) and centrifuged at 300 ×g for 10 minutes. Tumor-infiltrating lymphocytes (TILs) were isolated using density gradient centrifugation with Ficoll (GE Healthcare). The collected cells were washed with RPMI-1640 medium and assessed for cell counting using trypan blue.

### Single-cell RNA sequencing

Libraries were prepared using the Chromium controller following the protocol outlined in the 10x Chromium Next GEM Single Cell 5-v2 Cell Surface Protein User Guide (CG000330). Initially, cell suspensions were diluted with nuclease-free water to achieve a targeted cell count of 10,000. These suspensions were then mixed with master mix and loaded with Single Cell 5′ Gel Beads and Partitioning Oil into a Next GEM Chip K. Within the droplets, RNA transcripts from single cells were uniquely barcoded and reverse-transcribed.

The purified libraries were quantified using qPCR following the qPCR Quantification Protocol Guide (KAPA) and qualified using the Agilent Technologies 4200 TapeStation. Finally, the libraries were sequenced on the HiSeq platform (Illumina) according to the specified read length in the user guide.

### Single-cell RNA sequencing data collection and preprocessing

To ensure maximal hypothetical similarity to patient conditions as characterized in the TCGA study on esophageal cancer^5^, we filtered the HNSCC and GAC datasets accordingly. For the HNSCC dataset, we selected samples from patients without HPV infection and excluded samples collected from the oral cavity or tonsillar regions. For the GAC dataset, we chose samples from patients annotated as having chromatin instability (CIN) subtype from the original article.

For in-house and public EAC datasets with available raw sequencing files, we utilized CellRanger (v3.1.0)^41^ with CellRanger reference GRCh38 for quantifying raw expression counts for each sample. We then used filtered_feature_bc_matrix files from CellRanger outputs as count matrices. For initial quality control, we filtered cells with fewer than 200 unique genes, more than 4000 to 9000 unique genes (selected for each sample), or more than 15 or 25 percentages of mitochondrial gene (selected for each sample). Additionally, we filtered out cells identified as doublets with DoubletFinder^42^ with default parameters. For GAC datasets from Kumar et al.^13^, we utilized the given processed gene expression matrix file without further quality controls.

For each processed cohort, we conducted initial identification of epithelial cells with Seurat (v4.1.1)^43^. We first processed each cohort separately by normalizing datasets with log normalization, finding variable features, and filtering variable features that were ribosomal genes, TCR-related genes, immunoglobulin genes, or mitochondrial genes. With the filtered variable features, we scaled the normalized matrix and performed dimension reduction with PCA (30 PCs) followed by UMAP. We found clusters of the cells with resolution 0.4. Of the identified clusters, we used epithelial markers such as EPCAM, KRT19, KRT18, and CDH1 for separating epithelial cells from immune and stromal cells for separate integration steps. During this process, we also annotated major cell types such as T cells, B cells, myeloid cells, plasma cells, endothelial cells, and fibroblasts (**Supplementary Fig 1b**).

For immune and stromal cells, we first used SCTransform (v0.3.3)^44^ for normalization, scaling, and variable feature selection. We ran SCTransform separately for each sample and identified variable features within each cohort by integrating variable features from each sample with “SelectIntegrationFeatures’ function from Seurat. We then merged all samples for each cohort and used integrated variable features and SCTranform matrices for dimension reductions and clustering. Next, we tried to identify disassociation induced genes (DIGs) with an adapted method from Zheng et al.^45^ to minimize effects of genes upregulated during tissue dissociation. Briefly, we conducted high resolution (2.0) clustering, and used experimentally validated dissociation genes from Brink et al.^46^ for Fisher’s exact test with cluster markers. After adjusting p-values with Benjamini-Hochberg method, we selected dissociation-related clusters which had smaller than 0.005 of adjusted p-values for their marker genes. We then calculated frequencies of those marker genes across the dissociation-related clusters, and annotated genes that appeared for more than 40% of the dissociation-related clusters as *de novo* DIGs. We then combined *de novo* DIGs with the experimentally validated DIGs. To prevent any major cell type marker genes to be included for the combined DIGs, we calculated top 25 marker genes for the major cell types identified previously and excluded these genes from the combined DIGs. For any further processing, we excluded the DIGs from the variable gene lists.

After identifying the DIGs, we ran SCTranform for each patient again but excluded the DIGs from the variable genes this time. We then merged all samples from each cohort and used integrated variable features and SCTranform matrices for dimension reductions and clustering. During this, we used harmony on PC dimensions with cohort, sequencing platform, and patient for grouping variables, and used these harmony dimensions for further dimension reduction and clustering. We again used high resolution (2.0) clustering for each cohort and excluded any clusters with genes indicating low quality cells, such as mitochondrial genes or dominant ribosomal genes.

With the filtered samples, we attempted to match the gene symbols and excluded any unique genes from each cohort. For each cohort, we first converted all gene symbols to matched gene symbol version by using ‘alias2SymbolTable’ function from limma (v3.50.0)^47^. We then selected genes that exist for all four cohorts and removed any genes that did not pass such selection. After gene symbol matching, we reran per-sample SCTransform and merged all samples regardless of their cohorts for merged immune and stromal cells. With variable genes selected through ‘SelectIntegrationFeatures’ function, we carried out harmony-corrected dimension reductions and clustering with all immune and stromal cells. Again, with high resolution clustering, we removed low-quality cells for the last time. With the final filtered cells, we processed the integrated datasets with per-sample SCTransform and harmony-corrected dimension reduction to generate integrated immune and stromal cells. We confirmed fine distributions of cells from each dataset (**Supplementary Fig 1c**). With these cells, we annotated major cell types with corresponding marker genes.

For selected epithelial cells, we conducted several strict filtering steps for removal of any non-epithelial cells and identification of malignant epithelial cells. We first used the same method for gene symbol matching and merged all samples. Unlike immune and stromal cells, we did not use SCTransform for its tendency for over-correction of epithelial cells. After merging samples, we also used harmony-corrected dimension reduction but only ran harmony for correcting biases caused by sequencing platform. With high-resolution clustering, we removed any bad quality clusters or clusters with high immune cell markers. We also filtered out any non-epithelial cells annotated by SingleR (v1.6.1)^48^ HPCA (details described below). For identification of malignant epithelial cells, we used SCEVAN (v1.0.1)^49^ (details described below). After filtering non-malignant epithelial cells, we conducted the same harmony-corrected dimension reduction (only for sequencing platform) to acquire integrated malignant epithelial cells. After integration, patient heterogeneity was partially maintained especially for squamous carcinomas (**Supplementary Fig 1d**) while cells from different sequencing platforms were well mixed (**Supplementary Fig 1e**).

### TCGA bulk RNA sequencing data analysis

We acquired bulk-RNA sequencing datasets from TCGA with Esophageal Carcinoma (ESCA), Head-Neck Squamous Cell Carcinoma (HNSC), and Stomach Adenocarcinoma (STAD) studies with TCGAbiolinks (v2.22.1)^50^. Overall, we selected primary tumor samples for all studies. To match tumor characteristics to those of single cell datasets, we separated TCGA ESCA datasets^5^ into Esophageal Squamous Cell Carcinoma (ESCC) and Esophageal Adenocarinoma (EAC). For TCGA-HNSC^51^, we selected HPV negative samples only from larynx, hypopharynx, and oropharynx. For TCGA-STAD^52^, we choose samples with chromatin instability (CIN) subtype. For DEG analysis, we used ‘DESeq’ function from DESeq2 (v1.34.0)^53^ with variance stabilizing transformation (VST) for transformation of count data. We visualized z-scored expression of these DEGs for all four tumor types with pheatmap (v1.0.12)^54^. For dimension reduction with principal component analysis (PCA), we used ‘plotPCA’ function from DESeq2 with the transformed count matrix and visualized PC1 and PC2 coordinates of each sample with ggplot2 (v3.4.3)^55^.

### Pseudobulk analysis

To generate pseudobulk samples from single-cell RNA sequencing datasets, we either used all cells or subset specific cells (stromal and immune or malignant) and aggregated the raw count matrices for each patient with ‘aggregate.Matrix’ function from Matrix.utils (v0.9.8, https://github.com/cvarrichio/Matrix.utils). For DEG analysis and PCA following pseudobulk generation, we used the same approaches as for TCGA bulk sequencing datasets except for using regularized logarithm (rlog) for count data transformation considering smaller sample counts for pseudobulk samples.

### Reference-based automated annotation of epithelial cells

To identify epithelial cells, we utilized SingleR (v1.6.1)^48^ with Human Primary Cell Atlas (HPCA) reference. To strictly filter out non-epithelial cells, we only selected cells that were not annotated as one of the followings: B_cell’, ‘BM’, ‘BM & Prog.’, ‘CMP’, ‘DC’, ‘Endothelial_cells’, ‘Erythroblast’, ‘Gametocytes’, ‘GMP’, ‘HSC_-G-CSF’, ‘HSC_CD34+’, ‘Macrophage’, ‘MEP’, ‘Monocyte’, ‘MSC’, ‘Myelocyte’, ‘Neutrophils’, ‘NK_cell’, ‘Osteoblasts’, ‘Platelets’, ‘Pre-B_cell_CD34-’, ‘Pro-B_cell_CD34+’, ‘Pro-Myelocyte’, ‘T_cells’, ‘Fibroblasts’,’Smooth_muscle_cells.’

### Identification of malignant epithelial cells

To identify malignant cells from epithelial cells, we used SCEVAN (v1.0.1)^49^ for each patient sample. We combined all cells from integrated immune and stromal cells and integrated epithelial cells and separated those cells by their patient identities. For running SCEVAN for each patient sample, we used pipelineCNA function from SCEVAN with non-epithelial cells as normal cells. After running SCEVAN for each patient, we aggregated all results for epithelial cells and identified malignant epithelial cells.

### NMF-based metaprogram (MP) generation

First, we generated NMF modules for each sample with cNMF (v1.4.0)^56^. We prepared raw gene count matrix from each patient by separating integrated datasets by patient identities and filtering out genes that were not expressed by any cells for that sample. We ran cNMF with k values from 2 to 15 with 100 iterations and 4000 variable genes for malignant cells and 2000 variable genes for immune and stromal cells. We manually selected the optimal k value for each patient by evaluating k_selection_plot for the best balance between stability and error values. We re-ran the last cNMF ‘consensus’ step with the selected k values.

After running cNMF for each sample, we collected all NMF modules from all samples. For each patient, we ranked genes in each module with their calculated z-scores and ranked genes across the modules with their calculated TPM values. For each module, we selected genes with the ranks first across the modules and with the ranks that are above average within the module. We additionally filtered out any non-coding genes according to Consensus CDS database (assembly GRCh38.p14) and filtered out any non-variable genes. We then selected top 100 genes that satisfied all these conditions for each module. Using all modules with selected genes, we calculated jaccard similarities among all modules. We then filtered out any modules that did not have jaccard similarities above certain threshold with certain number of other modules. We selected jaccard similarity of 0.1∼0.2 with 3∼5 other modules depending on the optimal threshold for each major celltype.

With filtered modules, we generated co-occurrence network with the genes in the modules. We first filtered out genes that exist in fewer than 3 modules. After gene filtering, we made adjacency matrix with all remaining genes. We filtered out gene pairs with fewer than 3 co-occurrence within the modules. For each gene pair passing the threshold, we calculated their weighted co-occurrence count by dividing co-occurrence count with sum of gene appearance count for all modules. With the resulting adjacency matrix with weighted co-occurrence counts, we generated a weighted undirected gene network with ‘graph_from_adjacency_matrix’ function from igraph (v1.3.0)^57^.

With the gene network, we clustered genes with ‘cluster_infomap’ function with 100 number of trials. For each cluster, we ranked genes based on their degree centralities in the network. Of all gene clusters, we selected the clusters with at least 25 genes and chose maximum of 100 genes for each selected cluster. These final clusters were named MPs. We annotated each MP for its function based on pathway enrichment analysis and similarities to other public MPs. Finally, for malignant cells, MPs dominant with immune cell markers, mitochondrial genes, and ribosomal genes were filtered as noise MPs. For immune and stromal cells, MPs dominant with other immune cell markers, mitochondrial genes, and ribosomal genes were filtered as noise MPs.

### Scoring each cell with meta programs

For each cell, we used AUCell (v1.16.0)^58^ for evaluating whether the cell was ‘on’ or ‘off’ for the annotated meta programs. We calculated the cells’ AUC values with ‘AUCell_calcAUC’ with default parameters and manually selected the optimal AUC thresholds for each meta program. The cells passing the thresholds would be annotated to have the meta program ‘on.’ For analyzing proportions of ‘on’ cells for the meta programs, we calculated ratio of ‘on’ cells for each meta program to total cell counts for the cell type origin of the meta program for each patient. We then either used mean values of the proportions for tumor type-based analysis or calculated correlations of MP proportions among patients.

### Patient similarity dendrogram

To cluster patients with their similarities based on their malignant epithelial cells, we adapted ‘dendrogram’ function from scanpy^59^ to be usable in R. We averaged PC coordinates of the cells (25 for malignant cells) for each patient and calculated pearson correlations with these averaged PC coordinates among patients. This calculated correlation matrix was visualized with pheatmap (v1.0.12).

### Cell type composition analysis

For compositional analysis of cell subtypes identified after sub clustering analysis, We calculated proportions of each cell subtypes for each patient and group the proportions by tumor types. We then calculated p-values for the significance of compositional enrichment of cell subtypes by utilizing two-sided Wilcoxon rank sum tests for all combinations of tumor types. Only the significant p-values were indicated with asterisks (\**P* < 0.05; \*\**P* < 0.01; \*\*\**P* < 0.001).

### T cell annotation with projectTIL

For reference-based cell type annotation for CD4+ T cells and CD8+ T cells, we utilized ProjecTILs(v3.1.0)^24^ and their built-in T cell references (CD4T_human_ref_v1, CD8T_human_ref_v1). We separated cells from each tumor type and ran projecTILs for each tumor type without filtering any cells. After automated annotation, we calculated proportions of the annotated cell types for each tumor type and visualized with dot and line plots with ggplot2 (v3.4.0).

### Cellular trajectory analysis

For pseudotime trajectory analysis for CD8+ T cells, we used monocle3 (v1.2.9)^60^. We converted SCTransform matrix from Seurat object to monocle object using the default parameters. Using already calculated dimension reduction spaces from previous RNA processing steps, we learned principal graph of the monocle object with ‘learn_graph’ function from monocle3 without any prior information. We then ordered cells by choosing the root node closest to the naïve and memory cell populations by using ‘order_cells’ function. Because two diverging trajectories to exhausted T cells and HSP-high T cells were generated, we chose nodes in those cell populations as ending nodes with ‘choose_graph_segments’ function.

After generating two branches of developmental trajectories, we calculated signature scores along the pseudotime trajectories. The signature scores were calculated for each cell with ‘AddModuleScore’ function from Seurat (v4.1.1)^43^. We fitted the signature scores along the trajectories using ‘y ∼ bs(x, degree=3)’ formula in ‘geom_smooth’ function from splines (R-base package) and ggplot2 (v3.4.0). For calculation of transition proportion and ratio, we divided pseudotimes into early, intermediate, and late stages by bottom 33% and top 33% percentiles along the pseudotimes. For population proportions, we calculated divided number of cells in each stage of the trajectory for total number of cells of the trajectory for each patient. For transition ratio, we divided the number of cells in late stage by the number of cells in intermediate stage for each patient (Late / Intermediate ratio) or divided the number of cells in late stage by the number of cells in early stage for each patient (Late / Early ratio).

### Gene set enrichment analysis

For pathway enrichment analysis, we utilized enrichR (v3.2)^61^ with built-in databases. With mentioned databases for each analysis, we used adjusted p-values calculated with Fisher’s exact test and corrected for multiple hypotheses testing using Benjamini-Hochberg method. For visualization with bar plots, we took negative Log_10_ of adjusted p-values for the length of the bars, and the colors indicate overlap ratios that were calculated by dividing the number of input genes that are found in each term by the number of input genes. We chose 0.05 as significance threshold for the adjusted p-values.

### Single cell signature scoring & sources of the signature

Other than identifying ‘on’ and ‘off’ cells for meta programs, we scored signature scores for each cell by using ‘AddModuleScore’ from Seurat (v4.1.1) with default parameters except for ‘search = TRUE’ for searching synonyms for the input gene symbols. For T cell exhaustion signature, we used the following genes: *LAG3, HAVCR2, CTLA4, PDCD1, TIGIT, ENTPD1, ITGAE, TOX, CXCL13, KRT86*. For neo-antigen reactive signature, we combined genes from five neo-antigen reactive signatures. For stress responses from CD8^+^ T cells, we used stress response signature. All the gene signatures are listed in **Supplementary table 6.**

### Cell-cell interaction analysis using single-cell transcriptome data

To identify cell-cell interactions among different cell subpopulations, we used CellChat (v1.5.0)^40^ with the default interaction database. We selected log normalized RNA count matrices as input for CellChat and followed default pipelines for CCI inference. We visualized interactions among input cell types with circle plots with ‘netVisual_circle’ function from CellChat. We used weighted interaction strength that considered sizes of each cell type group.

### Gene signature scoring with bulk transcriptome data

To score enrichment of different gene signatures extracted from single-cell datasets for bulk RNA sequencing datasets, we used GSVA (v1.42.0)^62^ with default parameters. For direct comparison of signature scores between ICB responders and non-responders, we compared the calculated GSVA scores between two groups with two-sided Wilcoxon rank-sum test. Co-scores were calculated by averaging scores from two gene signatures. Diff-scores were calculated by subtracting a score from one gene signature by a score from another signature. For evaluating response predictive power of the signature scores, we used ‘roc’ function from pROC (v1.18.4)^63^ with gsva scores and response information (responder vs non-responder).

## Ethical approval and consent to participate

The studies were approved by the Institutional Review Board of Yonsei University Severance Hospital with IRB No 4-2016-0678. Written informed consent was obtained prior to enrollment and sample collection at Yonsei University Severance Hospital. The research conformed to the principles of the Helsinki Declaration.

## Availability of data and materials

The single-cell RNA sequencing data generated in this study is deposited in the Gene Expression Omnibus database and will be available to public upon publication. The remaining data are available within the Article or Supplementary Information.

## Authors’ contributions

S.Y.P., H.R.K., and I.L. conceived the study. S.B. performed single-cell transcriptome data analysis under the supervision of I.L. J.C. assisted bioinformatic analysis. S.Y.P. and H.R.K. organized clinical sample and data collections. M-H.H, Y.W.K, and D.K contributed to clinical sample collection. G.K. contributed sample preparation. M.H. provided advice on single-cell data analysis. I.L. and H.R.K. contributed to the financial and administrative support for this study. S.B., S.Y.P., H.R.K., and I.L. wrote the manuscript.

## Funding and Acknowledgement

This research was supported by the Bio & Medical Technology Development Program of the National Research Foundation funded by the Ministry of Science and ICT (2021R1A2C2094629, 2017M3A9E9072669 to H.R.K, 2022M3A9F3016364, 2022R1A2C1092062 to I.L.). The work was supported in part by Brain Korea 21(BK21) FOUR program. This work was supported by the Technology Innovation Program (20022947) funded by the Ministry of Trade Industry & Energy (MOTIE, Korea).

