## Supplementary Information for "Tumor microenvironment distinctions between esophageal cancer subtypes explain varied immunotherapy responses"

#### Contents

- ***Supplementary Tables 5, 13 (Supplementary Tables 1-4, 6-12, 14 are provided as tabular data files)***
- ***Supplementary References***
- ***Supplementary Figures***

#### Supplementary Tables

**Supplementary Table 1 |** Patient metadata

**Supplementary Table 2 |** Sample cell counts

**Supplementary Table 3 |** Marker genes for immune & stromal cell types

**Supplementary Table 4 |** Malignant and TME metaprograms

**Supplementary Table 5 |** Sources for gene signatures used in this study

| Reference | Gene signatures |
|---|---|
| Gavish et al., 2023 [1] | Cancer cell metaprograms |
| Barkley et al., 2022 [2] | Cancer cell metaprograms |
| Yuan et al., 2019 [3] | Cancer cell metaprograms |
| Curated | Exhaustion signature |
| Wischnewski et al., 2023[4] | Neo-antigen reactive signatures |
| Chu et al., 2023[5] | Stress response signature |
| Cabrita et al., 2020[6] | Tertiary lymphoid structure signature |
| Aran et al., 2017[7] | M1 Macrophage signature (Xcell) |
| Aran et al., 2017[7] | M2 Macrophage signature (Xcell) |
| Ye et al., 2019[8] | Hypoxia signature |
| Liberzon et al., 2015[9] | Interferon-gamma signature (MsigDB HALLMARK INTERFERON GAMMA RESPONSE) |

**Supplementary Table 6 |** Gene signatures used in this study

**Supplementary Table 7 |** Marker genes for $CD8^+$ T cell subsets

**Supplementary Table 8 |** Exhausted and Effector $CD8^+$ T cell exhaustion score and neo-antigen reactive score group DEGs

**Supplementary Table 9 |** Marker genes for $CD4^+$ T cell subsets

**Supplementary Table 10 |** Marker genes for B cell subsets

**Supplementary Table 11 |** Marker genes for Myeloid cell subsets

**Supplementary Table 12 |** TAM squamous cell carcinomas and adenocarcinoma DEGs

**Supplementary Table 13 |** Bulk RNA-seq samples with ICB treatment used in this study

| **Cohort** | **Cancer type** | **ICB treatment** |
|---|---|---|
| Chen et al., 2016[10] | Melanoma | Anti-CTLA4+Anti-PD1 |
| Kim et al., 2018[11] | Gastric Adenocarcinoma | Anti-PD1 |
| Lauss et al., 2017[12] | Melanoma | ACT |
| Gide et al., 2019[13] | Melanoma | (1) Anti-PD1 or (2) Anti-CTLA4 + Anti-PD1 |
| Rose et al., 2021[14] | Urothelial Carcinoma | Anti-PDL1 / Anti-PD1 |
| He et al., 2021[15] | Thymic Carcinoma | Anti-PD1 |
| Hsu et al., 2021[16] | Hepatocellular Carcinoma | Anti-PD1 & + Anti-CTLA4 / Anti-PD1 |

**Supplementary Table 14 |** Gene signatures from $CD8^+$T & TAM subsets

### Supplementary Figures

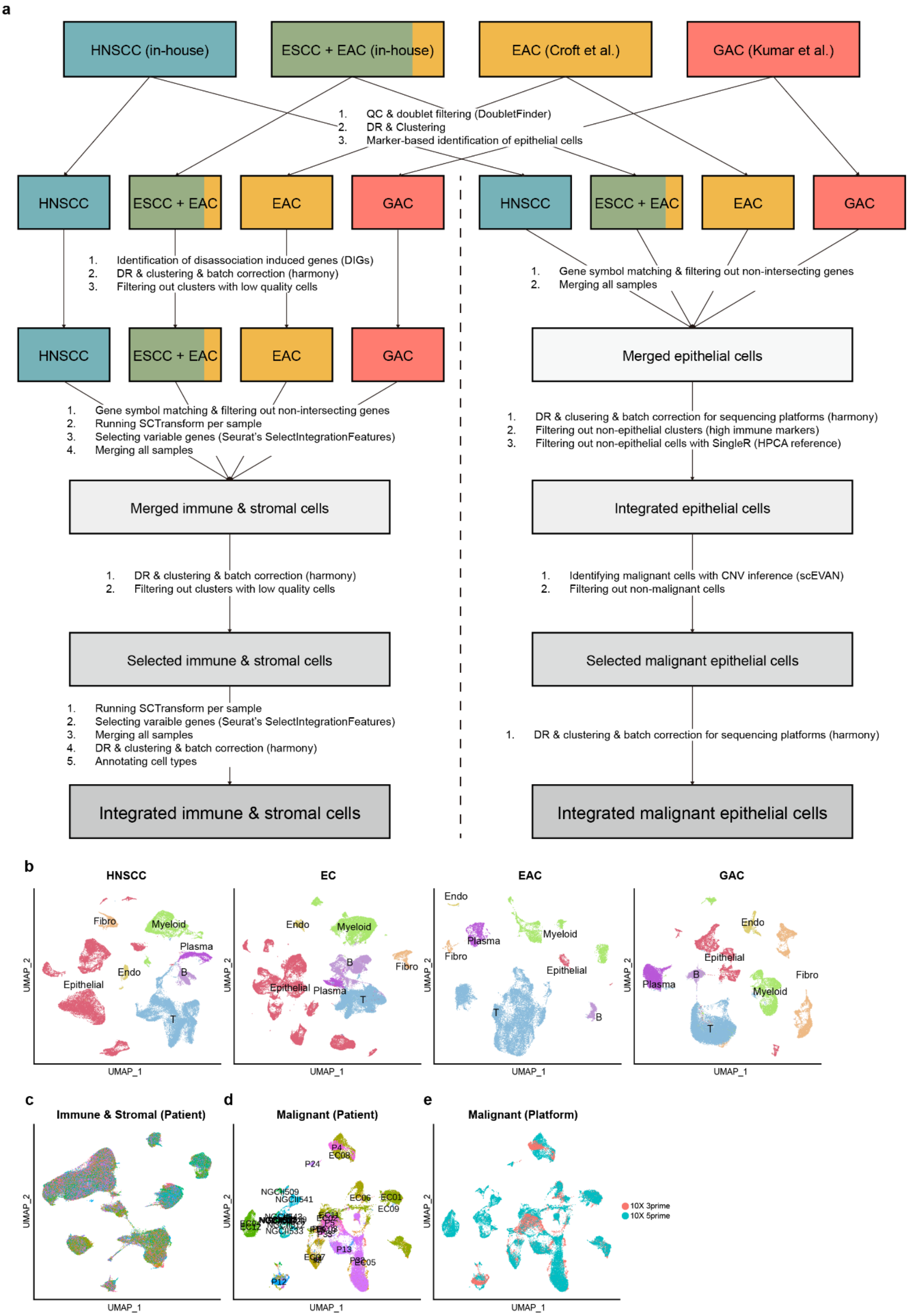

**Supplementary Fig. 1 | Overall scheme of single-cell RNA sequencing data preprocessing and integration. a,** Selected single-cell RNA-seq datasets from in-house cohorts and published studies and integration schemes for cells from those selected datasets. Each color indicates different cancer types. Used packages or functions are listed in the parenthesis. The Integration scheme for immune and stromal cells is on the left and the other integration scheme for epithelial cells is on the right. **b,** UMAP plot of total cells from each cohort. (First) In-house HNSCC cohort, (Second) In-house ESCC and EAC cohort, (Third) EAC cohort from Croft et al., Mol Cancer 2022, (Fourth) GAC cohort from Kumar et al., Cancer Discovery 2022. **c,** UMAP plot of integrated immune and stromal cells colored by their patient identities. **d,** UMAP plot of integrated malignant tumor cells colored by their patient identities. **e,** UMAP plot of integrated malignant tumor cells colored by sequencing platforms.

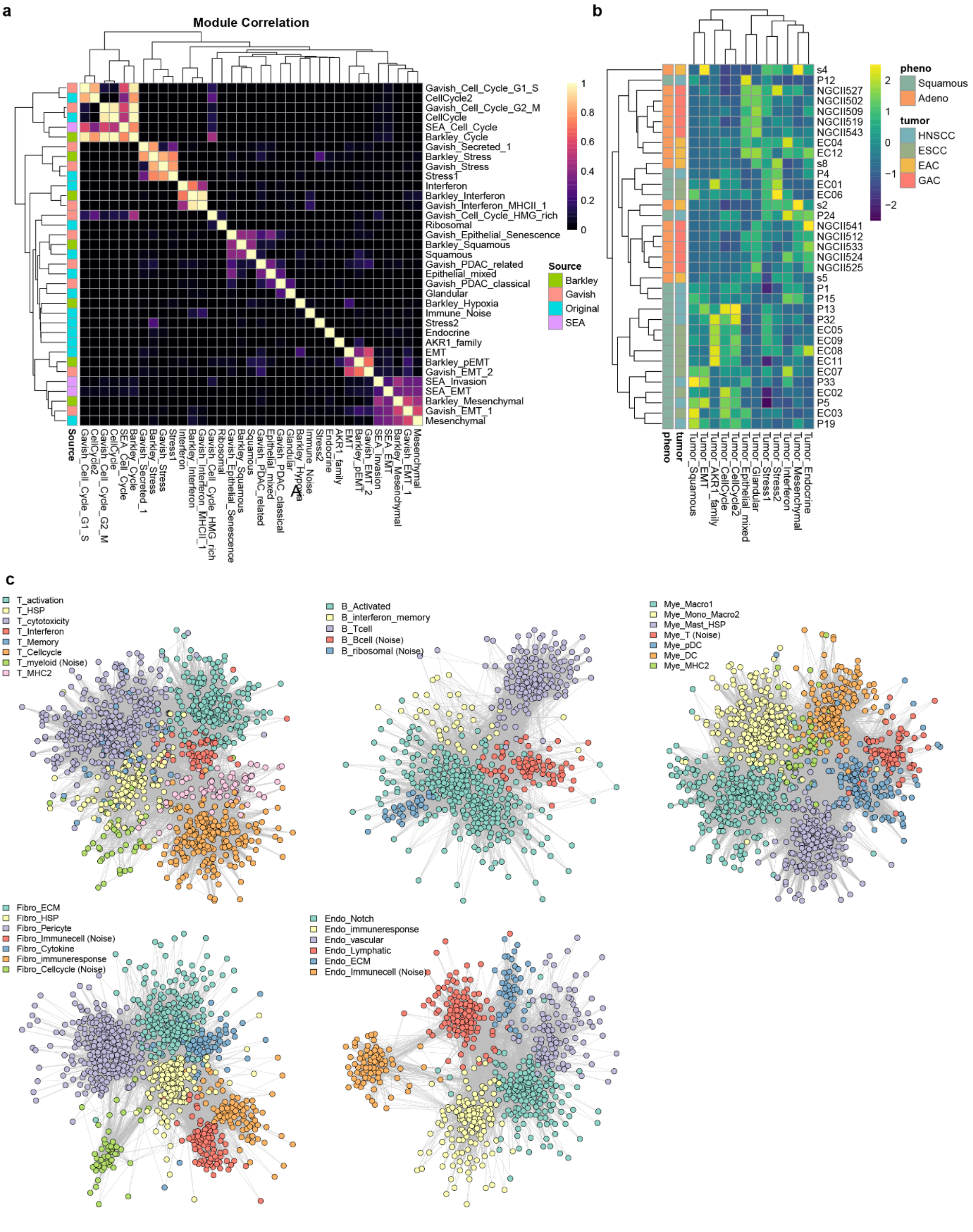


**Supplementary Fig. 2 | Malignant MPs and TME MPs. a,** Heatmap of jaccard similarities among the MPs from this study and three other public MPs. The rows were additionally colored by the sources of the MPs. The rows and the columns are clustered with hierarchical clustering. **b**, Heatmap for the proportions of 'on' cells from each patient for each MP. The rows and the columns are clustered with hierarchical clustering. The proportion values are column-wise z-scaled. **c,** Co-occurrence gene networks for MPs of each major cell type in TME. The nodes are the genes and are colored according to their MP classifications. The network layouts are in Fruchterman-Reingold layouts.

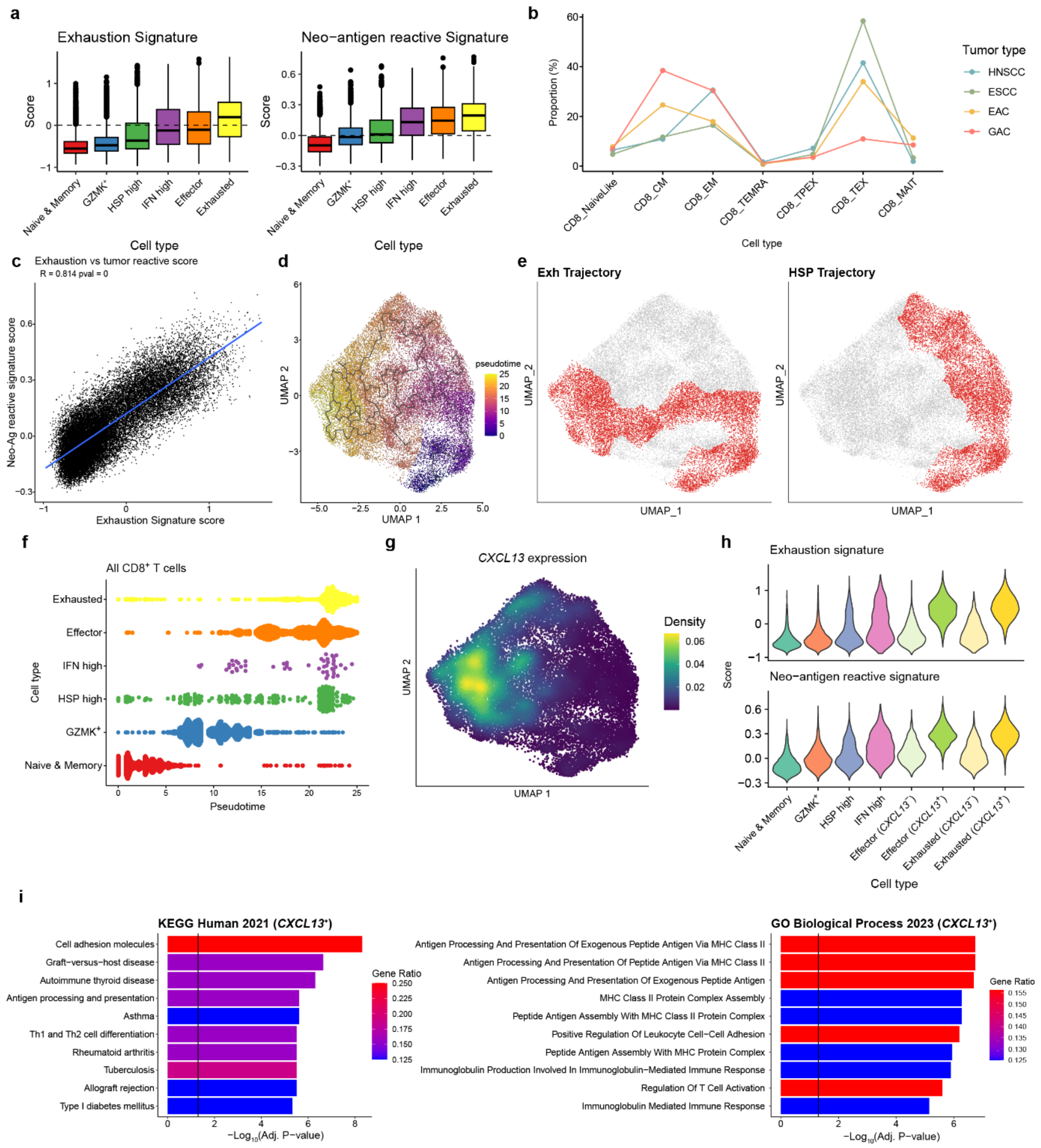

**Supplementary Fig. 3 | CD8$^+$T cells and subpopulations. a,** Boxplots of signature scores with (Left) exhaustion signature and (Right) neo-antigen reactive signature. The boxplots are grouped for each of CD8$^+$ T subpopulations. **b,** Proportions of subpopulations of CD8$^+$ T cells annotated with projecTIL. The proportions are calculated for each tumor type. **c,** Correlation between exhaustion scores and neo-antigen reactive scores for CD8$^+$ T cells. Pearson correlation coefficient and p-values are indicated. **d,** UMAP plots with all trajectory branches calculated from monocle 3. Colors indicate pseudotime. **e,** UMAP plots of CD8$^+$ T cells highlighted for cells involved in (Left) exhaustion trajectory and (Right) HSP trajectory. **f,** All cells aligned through the calculated pseudotimes grouped and colored by $CD8^+$ subpopulations. **g,** UMAP plot for density of *CXCL13* expressing $CD8^+$ T cells. **h,** Violin plots of signature scores of exhaustion signature (top) and neo-antigen reactive signature (bottom) for subpopulations of $CD8^+$ T cells. Effector and exhausted $CD8^+$ T cells were further separated by *CXCL13* expression. **i.** Pathway analysis with DEGs from comparing *CXCL13*$^+$ $CD8^+$ T cells to *CXCL13*$^-$ $CD8^+$ T cells. Colors of the bar indicate ratio of genes from the DEGs that are in gene list for each pathway term. Black vertical lines indicate $q$-value threshold of 0.05. KEGG Human (2021) and GO Biological Process (2023) databases were used.

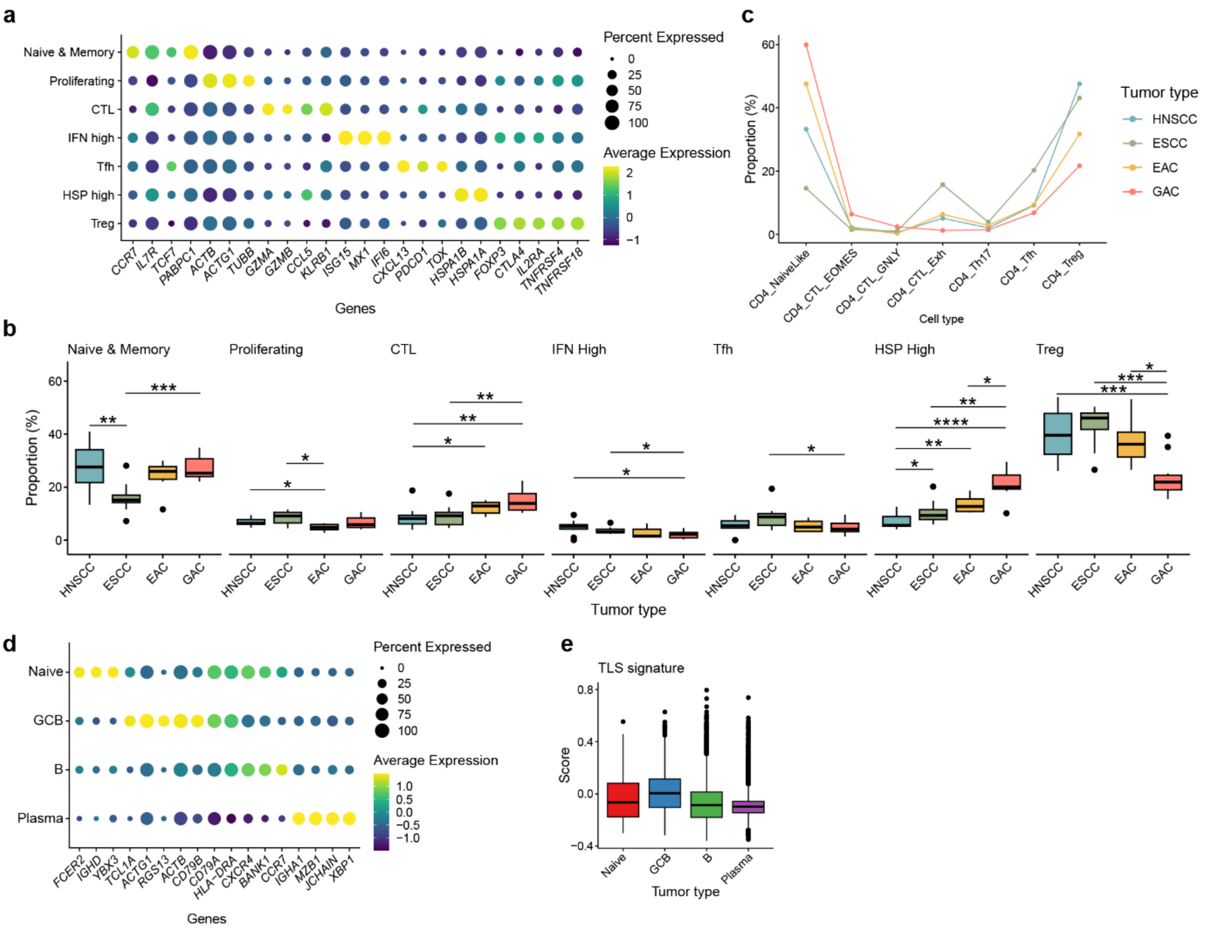


**Supplementary Fig. 4 | CD4$^+$ T cells and B cells. a,** Dotplot of markers genes for each subpopulation of CD4$^+$ T cells. The color indicates scaled average expression for each subpopulation and the sizes indicate percentage of cells that are expressing each gene for each subpopulation. **b,** Proportion of CD4$^+$ T cell subpopulations for each patient. The proportions are visualized by the boxplots for each cancer type and each CD4$^+$ T cell subpopulation. **c,** Proportions of subpopulations of CD4$^+$ T cells annotated with projecTIL. The proportions are calculated for each tumor type. **d,** Dotplot of markers genes for each subpopulation of B cells. The color indicates scaled average expression for each subpopulation and the sizes indicate percentage of cells that are expressing each gene for each subpopulation. **e,** Boxplots of signature scores of tertiary lymphoid structure signature from each cell. The boxplots are grouped for each B cell subpopulation.

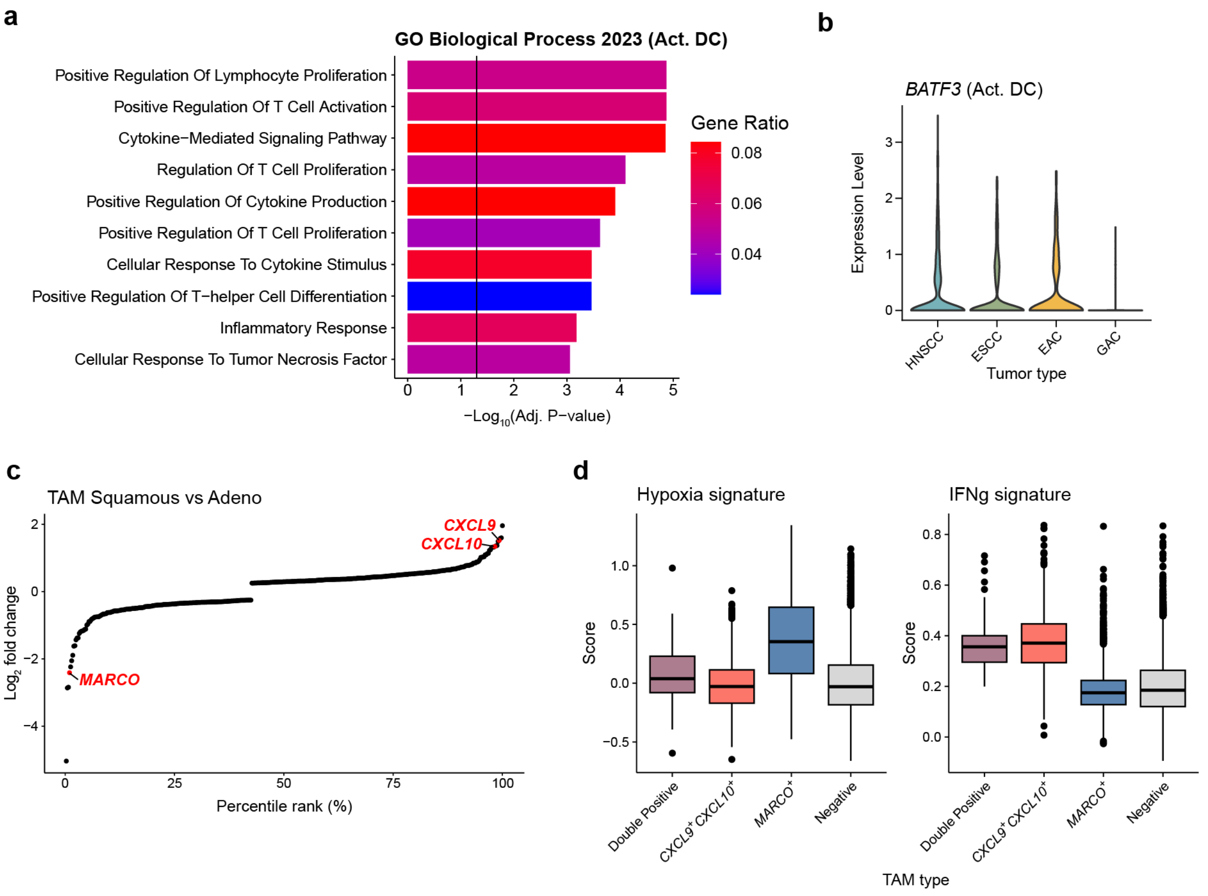


**Supplementary Fig. 5 | Myeloid cells. a,** Pathway enrichment analysis with activated DCs. Colors of the bar indicate ratio of genes from the DEGs that are in gene list for each pathway term. Black vertical lines indicate $q$-value threshold of 0.05. GO Biological Process (2023) database was used. **b,** Violin plots of expression of BATF3 for activated DCs from each tumor type. **c,** Differentially expressed genes between TAMs of SCC (ESCC and HNSCC) and AC (EAC and GAC) visualized with percentile ranks based on $\log_2$ fold change (x-axis) and $\log_2$ fold change (y-axis). **d,** Boxplots of signature scores for each TAM classification with hypoxia, Tumor inflammation, macrophage-specific tumor inflammation, and interferon gamma signatures.
